# Extant Sequence Reconstruction: The accuracy of ancestral sequence reconstructions evaluated by extant sequence cross-validation

**DOI:** 10.1101/2022.01.14.476414

**Authors:** Michael A. Sennett, Douglas L. Theobald

**Affiliations:** Brandeis University Department of Biochemistry Waltham, MA 02453 USA

## Abstract

Ancestral sequence reconstruction (ASR) is a phylogenetic method widely used to analyze the properties of ancient biomolecules and to elucidate mechanisms of molecular evolution. Despite its increasingly widespread application, the accuracy of ASR is currently unknown, as it is generally impossible to compare resurrected proteins to the true ancestors. Which evolutionary models are best for ASR? How accurate are the resulting inferences? Here we answer these questions using a cross-validation method to reconstruct each extant sequence in an alignment with ASR methodology, a method we term “extant sequence reconstruction” (ESR). We thus can evaluate the accuracy of ASR methodology by comparing ESR reconstructions to the corresponding known true sequences.

We find that a common measure of the quality of a reconstructed sequence, the average probability, is indeed a good estimate of the fraction of correct amino acids when the evolutionary model is accurate or overparameterized. However, the average probability is a poor measure for comparing reconstructions from different models, because, surprisingly, a more accurate phylogenetic model often results in reconstructions with lower probability. While better (more predictive) models may produce reconstructions with lower sequence identity to the true sequences, better models nevertheless produce reconstructions that are more biophysically similar to true ancestors. In addition, we find that a large fraction of sequences sampled from the reconstruction distribution may have fewer errors than the single most probable (SMP) sequence reconstruction, despite the fact that the SMP has the lowest expected error of all possible sequences. Our results emphasize the importance of model selection for ASR and the usefulness of sampling sequence reconstructions for analyzing ancestral protein properties. ESR is a powerful method for validating the evolutionary models used for ASR and can be applied in practice to any phylogenetic analysis of real biological sequences. Most significantly, ESR uses ASR methodology to provide a general method by which the biophysical properties of resurrected proteins can be compared to the properties of the true protein.

## Introduction

Pauling and Zuckerkandl (Pauling and Zuckerkandl 1963) first proposed that one could take a family of modern proteins, reconstruct the sequences of their ancestors, and “resurrect” the ancestral proteins by synthesizing them and studying their properties experimentally. Such ancestral sequence reconstruction (ASR) methods have now become widely used to analyze the properties of ancient biomolecules and to elucidate the mechanisms of molecular evolution. By recapitulating the structural, mechanistic, and functional changes of proteins during their evolution, ASR has been able to address many fundamental and challenging evolutionary questions, areas where more traditional methods have failed (Akanuma et al. 2013; Boucher et al. 2014; Clifton et al. 2018; Dean and Thornton 2007; Harms and Thornton 2010; Hochberg and Thornton 2017; Kaltenbach et al. 2018; Liberles et al. 2012; Nguyen et al. 2017; Pillai et al. 2020). Furthermore, ASR methodology has been highly successful in addressing biophysical problems of modern proteins (Nicoll et al. 2023), such as unraveling the mechanism of cancer drug specificity in a modern kinase family (Wilson et al. 2015), teasing apart cryptic epistatic effects within a hormone receptor (Ortlund et al. 2007), clarifying puzzling sequence-structure-function relationships in enzymes (Wouters et al. 2003), and explaining why certain features of modern proteins are not functionally optimized (Finnigan et al. 2012) (these and many other examples reviewed in Hochberg and Thornton (2017),Thomson et al. (2022), and (Dube et al. 2022)). Despite the tangible successes of ASR, the accuracy of its reconstructions is still currently unknown, because it is generally impossible to compare resurrected proteins to the true ancient ancestors.

Ancestral resurrected proteins often possess remarkable physical properties that are absent in their modern counterparts, such as high thermostability, catalytic versatility, and resistance to damage (Risso et al. 2017; Risso et al. 2018; Spence et al. 2021; Trudeau et al. 2016; Zakas et al. 2017). However, these exceptional properties might be artifacts resulting from the known biases in the protein reconstruction process rather than genuine characteristics of the ancestral proteins. Such potential biases could affect any global property of a protein (Akanuma 2017; Krishnan et al. 2004; Matsumoto et al. 2015; Risso et al. 2018; Thornton 2004; Trudeau et al. 2016; Wheeler et al. 2016; Williams et al. 2006; Yang 2006).

The most widely used ASR methods employ model-based probabilistic inference, such as maximum likelihood (ML) and Bayesian methodology. Hence, accurate ancestral reconstructions rely on accurate phylogenetic models. The most common model in molecular evolution is a time-reversible Markov model of residue substitution that assumes independent sites, rate variation among sites, a global equilibrium frequency distribution, and is homogeneous across sites and throughout the phylogeny (Felsenstein 1981; Yang 1994). Many other more biologically realistic evolutionary models have been proposed that relax various combinations of these model assumptions. Despite this rich theoretical framework, we presently have very limited methods for assessing the adequacy of the evolutionary models used for ASR.

Model-based probabilistic ASR methods predict a distribution of states for a reconstructed ancestor rather than a single sequence. The combinatorial number of plausible ancestral states is often astronomically high, and it is generally impossible to study them exhaustively. In practice, this problem is simplified by resurrecting only the single most probable (SMP) ancestral sequence as a proxy for what may have occurred in the past (Chang et al. 2002; Gaucher et al. 2003; Thornton et al. 2003). One proposed justification for using the SMP sequence is that it is expected to have the fewest errors relative to the true sequence (Eick et al. 2017), but this hypothesis has yet to be verified using real biological sequences. Although it is intuitively reasonable to focus on the SMP sequence, the amino acid composition of the SMP is known to be systematically biased, which can lead to downstream biases in experimental structure-function studies that depend on the SMP sequence (Krishnan et al. 2004; Williams et al. 2006).

The most direct method for validating ASR is to compare the reconstructed ancestral protein to the true ancestral protein. Such comparisons have been performed using proteins from directed evolution and simulated data with known alignments and phylogenies. Computational studies have focused on simulated data from approximate models of evolution that rely on various simplifying assumptions (Williams et al. 2006; Zhang and Nei 1997). Experimental studies have largely focused on reconstructing ancestral sequences from directed evolution experiments that are limited in their sequence divergence and relevance to natural evolution (Randall et al. 2016). It is unclear whether the results generalize to real biological systems with unknown phylogenies and uncertain alignments of sequences that have evolved on a geological timescale (Randall et al. 2016; Williams et al. 2006).

Here we propose a method, which we call Extant Sequence Reconstruction (ESR), that can assess the accuracy of ASR methodology by comparing reconstructed sequences to the corresponding true proteins. With time-reversible evolutionary models there is no distinction between ancestor and descendant. ESR uses this well-known property to effectively invert the traditional ASR calculation, using standard ASR methodology to reconstruct a modern protein sequence. A reconstruction of an extant protein provides both an SMP sequence and a reconstructed sequence distribution, just as with reconstructions of ancestral nodes. Because extant reconstructions are calculated in the same way as ancestral reconstructions—using the same probabilistic methodology, phylogeny, alignment, and evolutionary model—extant reconstructions should largely share the same accuracies, limitations, biases, and statistical characteristics as ancestral reconstructions. With an extant reconstruction we know the true sequence and thus can validate our prediction by direct comparison with truth, thereby providing a direct test of ASR methodology.

Using ESR on multiple sequence datasets and proteins, we quantify the accuracy of reconstructions by comparison to their corresponding true sequences. Our results highlight the critical importance of model selection for determining the best evolutionary model for ASR. We find that a common measure of the quality of a reconstructed SMP sequence, the average probability of the sequence, is indeed a good estimate of the fraction of the sequence that is correct when the evolutionary model is accurate or overparameterized. However, we also find that the average probability of the SMP reconstruction is a poor measure for comparing different SMP reconstructions, because more accurate phylogenetic models typically result in SMP reconstructions with lower probability and fewer correct residues. Though this result may initially appear paradoxical, we show that it is an expected feature of more realistic phylogenetic models that are not optimizing the fraction of correct ancestral amino acids. Rather, better evolutionary models will often opt to make more biophysically conservative mistakes rather than make fewer non-conservative mistakes. Our results suggest that a more reliable indicator of the quality of a reconstruction is the entropy of the reconstructed distribution, which provides an estimate of the log-probability of the true sequence. ESR is a widely applicable method for validating ASR evolutionary models and predictions. ESR can be used to in practice to evaluate any phylogenetic analysis of real biological sequences. While we have focused here on sequence-based characterization of ESR predictions, this work provides the foundation for future work in which we will use ESR to experimentally assess the accuracy of ASR methods by comparing the biophysical properties of reconstructed and resurrected proteins to the true proteins.

## Results

### Simulations show model misspecification can result in biased ASR probabilities

An important statistic that is commonly used to gauge the quality of an ancestral reconstructed sequence is its *average probability*, defined as the average over sites of the probabilities of the amino acids in the sequence (Equation 8 in *Methods*). The average probability of a sequence is equal to the expected fraction of correct amino acids in the sequence. Hence, experimentalists typically choose the SMP sequence to resurrect in the lab, because the SMP sequence has the fewest expected number of errors of all possible reconstructed sequences (Eick et al. 2017). Of course, this will only be valid if the reconstruction probabilities for the SMP sequence accurately reflect uncertainty in the amino acid state, which in turn depends on how well the evolutionary model describes the true biological process that generated the sequence data. It has been claimed, for instance, that overly simple models will give inaccurate reconstruction probabilities (Matsumoto et al. 2015; Songyang et al. 1995). A misspecified model might over-optimistically produce reconstruction probabilities that are systematically too high (if it is too simple and fails to capture relevant biological features), or the model might pessimistically under-estimate the reconstruction probabilities (perhaps if it is overfit).

Hence, an important open question in ASR is whether the amino acid probabilities in an ancestral distribution accurately reflect our uncertainties in the character states. We can quantify the accuracy of reconstruction probabilities by considering the sites in an SMP sequence as a series of independent Bernoulli trials. Each SMP amino acid selected has a probability of success. Reconstructed sites that have, say, 80% probability should actually be correct 80% of the time, on average. Consequently, if the average amino acid probability for an SMP sequence is 95% and our probabilities are accurate, then we should expect that roughly 95% of the predicted amino acids in the SMP sequence are indeed correct when compared to the true sequence, within counting error.

To evaluate the accuracy of reconstruction probabilities we need to compare ancestral reconstructions to the true ancestral sequences and calculate the fraction of correct amino acids, but we rarely know the true ancestral sequences for real biological datasets. One solution to this problem is to use simulated data in which we know the true ancestral sequences. Therefore, as an initial analysis, we simulated ancestral and extant sequences along the ML phylogenies for L/MDH, Abl/Src-kinase, and terpene synthases using the corresponding ML estimates from the LG+FO+G12 model of evolution obtained from our real protein datasets. Simulations and analyses were replicated ten times. Using these simulated datasets, we ascertained how the probabilities of a reconstructed ancestral SMP sequence are affected by evolutionary models of increasing complexity. We performed ASR for all simulated datasets by fixing the true tree topology while inferring ancestors, branch lengths, and model parameters under various models of evolution. In addition to using correctly specified models of evolution with various levels of parameterization, we also explore models that are intentionally misspecified.

For misspecified models (i.e., models with an incorrect functional form or constants set to incorrect values relative to the true model), the average probability of a reconstructed ancestral sequence overestimates the fraction correct and results in fewer correct residues (Fig. 1a-b, Supplementary Figures S1a, S2a, and S3a-c). However, for the correctly specified LG and parameter-rich GTR20 model the average SMP sequence probability is an accurate estimate of fraction correct. This result is robust to errors in branch length estimation, since fixing the branch lengths to their true values results in similar overly optimistic reconstruction probabilities, at least in trees of non-trivial number of taxa (Supplementary Figures S4 and S5).

**Fig. 1:**
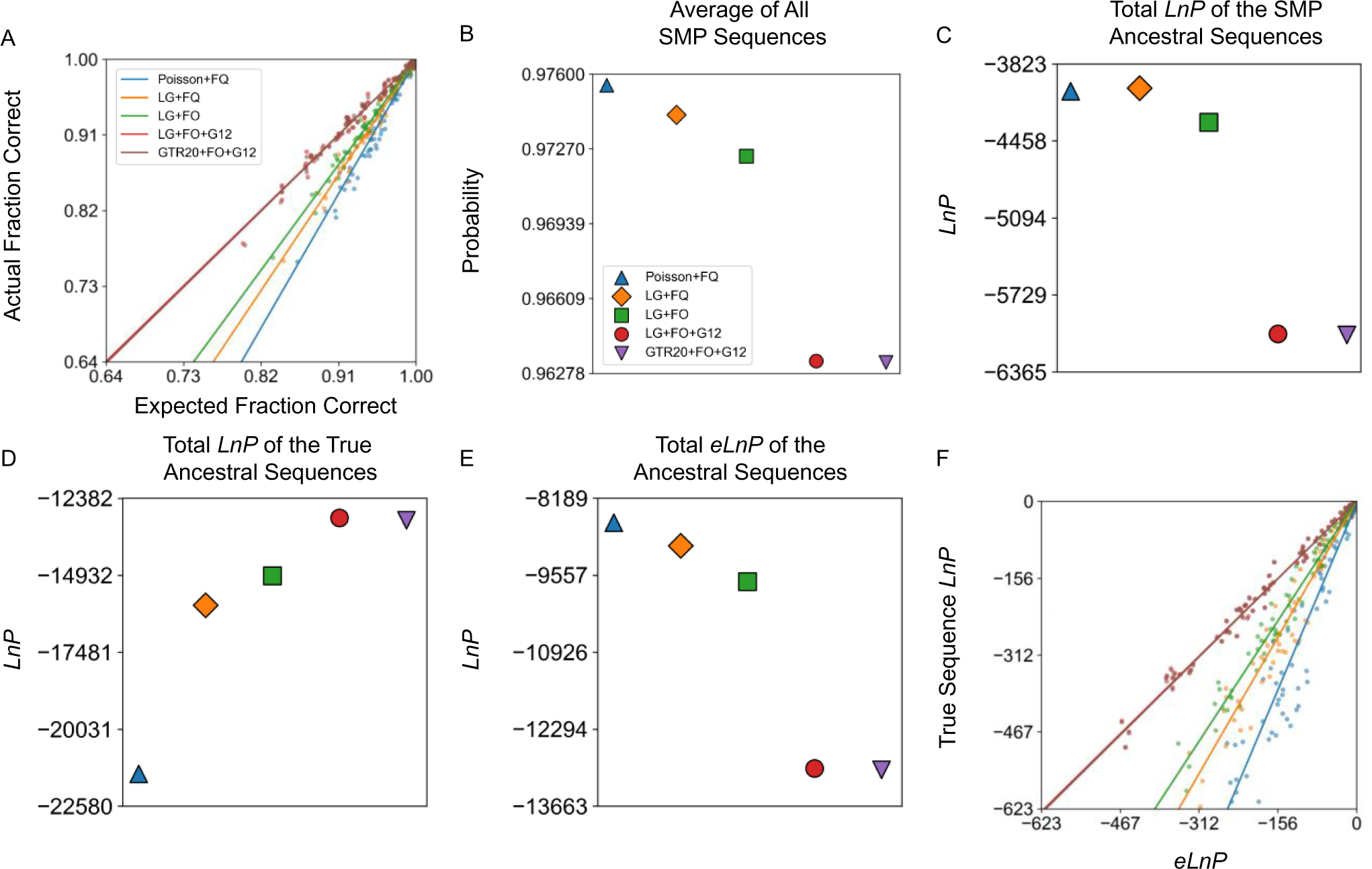
ASR probabilities are accurate when the model is true or overparameterized for LG+FO+G12 simulated sequences. Ten sets of ancestral sequences were simulated using the LG+FO+G12 model of evolution using an experimental L/MDH phylogeny (inferred from real data using LG+FO+G12). Analyses of ancestral reconstructions for the tenth dataset are shown. (*a*) A plot of the actual fraction correct against expected fraction correct for each reconstructed ancestral sequence in a simulation for each model of evolution. The corresponding line of best fit is shown for each model. (*b*) The average of all average SMP sequence probabilities for each model of evolution. (*c-e*) The total *LnP* of all SMP sequences, true sequences, and *eLnP* for each model of evolution. (*f*) True sequence *LnP* plotted against *eLnP* for each reconstructed ancestral sequence for each model of evolution. The slopes for (*a*) and (*f*) are given in Supplementary Table 2. The values for (*c*-*e*) are given in Supplementary Table 8.

To further explore the effects of misspecification on the accuracy of ancestral reconstruction probabilities, we also simulated ancestral and extant sequences using one of the simplest possible evolutionary models, the Poisson+FQ model, along the L/MDH, Abl/Src-kinase, and terpene synthase ML tree topology. For data simulated with this simple model, we find that the bias in probability estimation is lower in magnitude than with the LG+FO+G12 simulations (Fig. 2a-b, Supplementary Figures S6a and S7a).

**Fig. 2:**
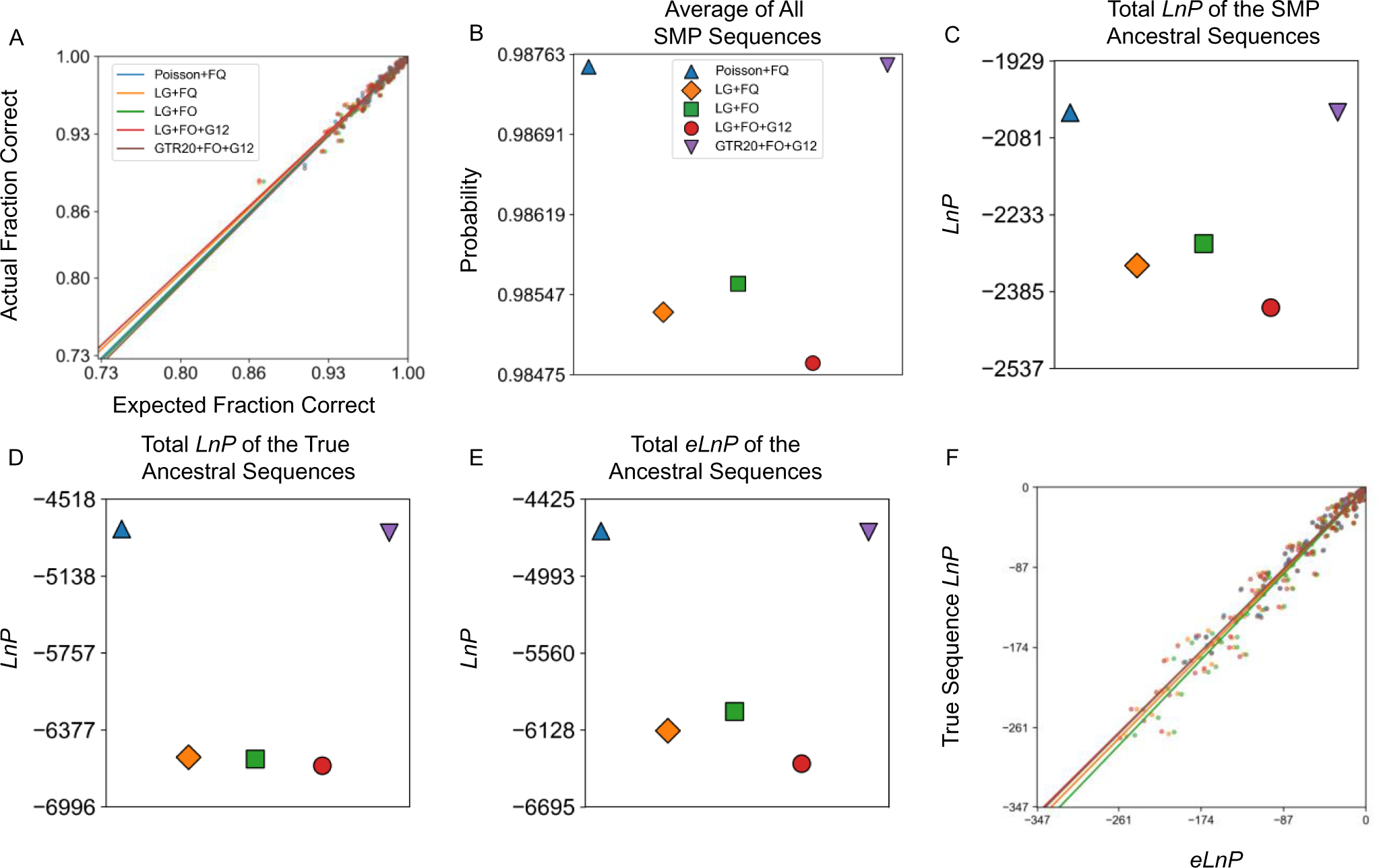
ASR probabilities for L/MDH Poisson+FQ simulated sequences are accurate even when the model is misspecified. Panels are the same as Fig. 2, except ten sets of ancestral sequences were simulated using the Poisson+FQ model of evolution on an experimental L/MDH phylogeny inferred using Poisson+FQ model. The slopes for (*a*) and (*f*) are given in Supplementary Table 3. The values for (*c*-*e*) are given in Supplementary Table 10.

For every model, a linear regression of actual fraction correct on expected fraction correct (i.e., average probability) gives positive slopes. As a result, for a given phylogenetic analysis with a specific evolutionary model, sequences with higher average probability are expected to have a higher actual fraction correct.

### More accurate models decrease the probability of the ancestral SMP sequence

Models that are most able to predict unobserved data will include parameters that correctly approximate some underlying process that generated the data. Hence, we hypothesized that more realistic models (*i*.*e*., those that take into account some underlying feature of data generation) should improve the accuracy of reconstructed distributions for ancestral sequences. For example, it is reasonable to expect that SMP sequences constructed using better models should have a higher overall probability, higher average amino acid probabilities, and by extension fewer amino acid errors with respect to the true sequences. In particular, for ancestral sequences generated using the LG+FO+G12 model, we might expect that the naïve Poisson model should produce less probable ancestral SMP sequences, while the LG+FO+G12 model should produce more probable ancestral SMP sequences.

We tested these expectations by comparing the true ancestral sequences to the corresponding SMP ancestral reconstructions from various models of evolution. For our datasets simulated under the LG+FO+G12 model of evolution, we calculated the overall average probability of all SMP ancestral sequences reconstructed for each model of evolution (Fig. 1b, Supplementary Figures. S1b and S2b). Surprisingly, as the model includes more parameters that help explain the underlying generative model for the data, the overall average probability of an SMP ancestral reconstruction decreases, resulting in more expected amino acid errors in the ancestral SMP sequence.

The exact probability of the SMP sequence can be calculated by taking the product over sites of the probability of each amino acid in the SMP sequence (Equation 6 in Methods). Because these probabilities are extremely small in general, it is conventional to take the natural logarithm of the SMP ancestral sequence probability (*LnP*). We calculated the total *LnP* for all SMP ancestral sequences in a phylogeny for each model of evolution (Fig. 1c, Supplementary Figures. S1c and S2c). Like the average probability, the total *LnP* for SMP ancestral sequences decreases as the model becomes more realistic.

### More accurate models increase the probability of the true ancestral sequence

Why do more predictive models result in reconstructed SMP sequences with lower probabilities? At each site, the probabilities associated with each amino acid must sum to 1. If the probability of the SMP residue decreases, then the probability of at least one other amino acid must increase. We then might expect a better model to improve the probability of the true residue when the SMP residue is incorrect. To see if this occurs, we calculated the total *LnP* of the true ancestral sequence for each model of evolution (Fig. 1d, Supplementary Figures. S1d and S2d). We see that the total *LnP* of the true sequences improve as the model is closer to truth or overparameterized. Hence, a more predictive model improves the probability of predicting the true ancestral sequence at the expense of the SMP sequence.

### The expected *LnP* is an accurate estimate of true ancestral sequence *LnP* for correctly specified models

The SMP sequence is known to have biased amino acid residue propensities, and hence any global property of the SMP sequence may also display a corresponding bias (Krishnan et al. 2004; Matsumoto et al. 2015; Williams et al. 2006). The reconstructed ancestral distribution contains all the information that the evolutionary model can provide about the true, unobserved ancestral sequence. Rather than analyzing the SMP sequence, a more statistically sound method uses the reconstructed ancestral distribution to calculate the expected value for properties of the unobserved ancestral sequence (Matsumoto et al. 2015).

For example, the *LnP* of the true ancestral sequence is unknown for real biological data, but it can be consistently estimated by the expected *LnP* (*eLnP*) if the model is correctly specified. Importantly, the *eLnP* can be calculated from the reconstructed distribution without knowing the identity of the true sequence (eqn. 7). The *eLnP* is mathematically equivalent to the negative of the entropy of the reconstructed distribution, which serves as a measure of the overall uncertainty in the distribution in specifying the reconstructed sequence.

To determine if the *eLnP* is an accurate estimate of the true ancestral sequence *LnP* we thus calculated the total *eLnP* for different protein families and models of evolution (Fig. 1e, Supplementary Figures. S1e and S2e). Like the total *LnP* of ancestral SMP sequences, the total *eLnP* decreases as the model more closely reflects the true generating model. Interpreting the *eLnP* as the negative entropy, we see that misspecified models assign a lower uncertainty to their predictions of the ancestral states.

Since the *eLnP* is a statistical estimate of the *LnP* of the true sequence, from our LG+FO+G12 simulations we can compare the calculated *eLnP* to the true sequence *LnP* for the different models of evolution (Fig. 1f, Supplementary Figures S1f and S2f). When the model is correctly specified, the *eLnP* is an accurate estimate of the true sequence *LnP*; otherwise, the *eLnP* overestimates the *LnP* of the true sequence. This behavior is similar to that of the SMP average probability, which also overestimates the fraction of correct residues with poor models.

### Extant Sequence Reconstruction (ESR): Cross-validation produces a reconstructed probability distribution for observed modern sequences

Our simulations have shown that ancestral sequence reconstruction probabilities are accurate when the model is correctly specified. The sequences generated from these simulations assume a time-reversible Markov model of residue substitution, site independence, global equilibrium frequencies, and homogeneity across sites and time. However, these assumptions may not hold for the underlying processes that generate real biological sequences, and hence it may not be valid to generalize from simulated data to real biological sequences. Furthermore, it would be of practical benefit to be able to apply our analyses to real experimental sequence data, rather than being limited to artificial simulations. To address these concerns, in the following we extend ASR methodology to the reconstruction of real modern protein sequences using a site-wise cross-validation (CV) method.

Similar to how ASR produces a reconstructed probability distribution for internal ancestral nodes of a tree, site-wise CV produces a reconstructed probability distribution for extant sequences found at the terminal nodes of a tree. When using time-reversible models, like all models considered in this paper, ancestral nodes are modeled identically as leaf nodes (*e.g.*, any leaf node may be treated as the ancestral root of the tree). Site-wise CV produces the conditional probability distribution of a modern site using the ML parameters from a model that has not seen the true modern amino acid at that site, thereby removing any circularity in the reconstruction. For example, to reconstruct the site for extant sequence *A*, we omit the residue at that site from the alignment and use the resulting training alignment to find the ML estimates of the tree and model parameters (Fig. 3, right top). With those ML parameters, we then calculate the probabilities of all 20 possible amino acids at that extant site using the same method used in conventional ASR (see equations (4) and (5) in the Methods):

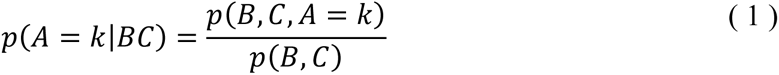

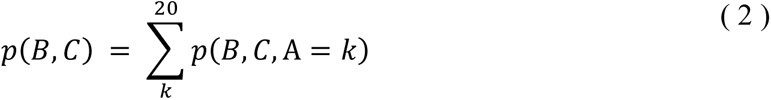

**Fig. 3:**
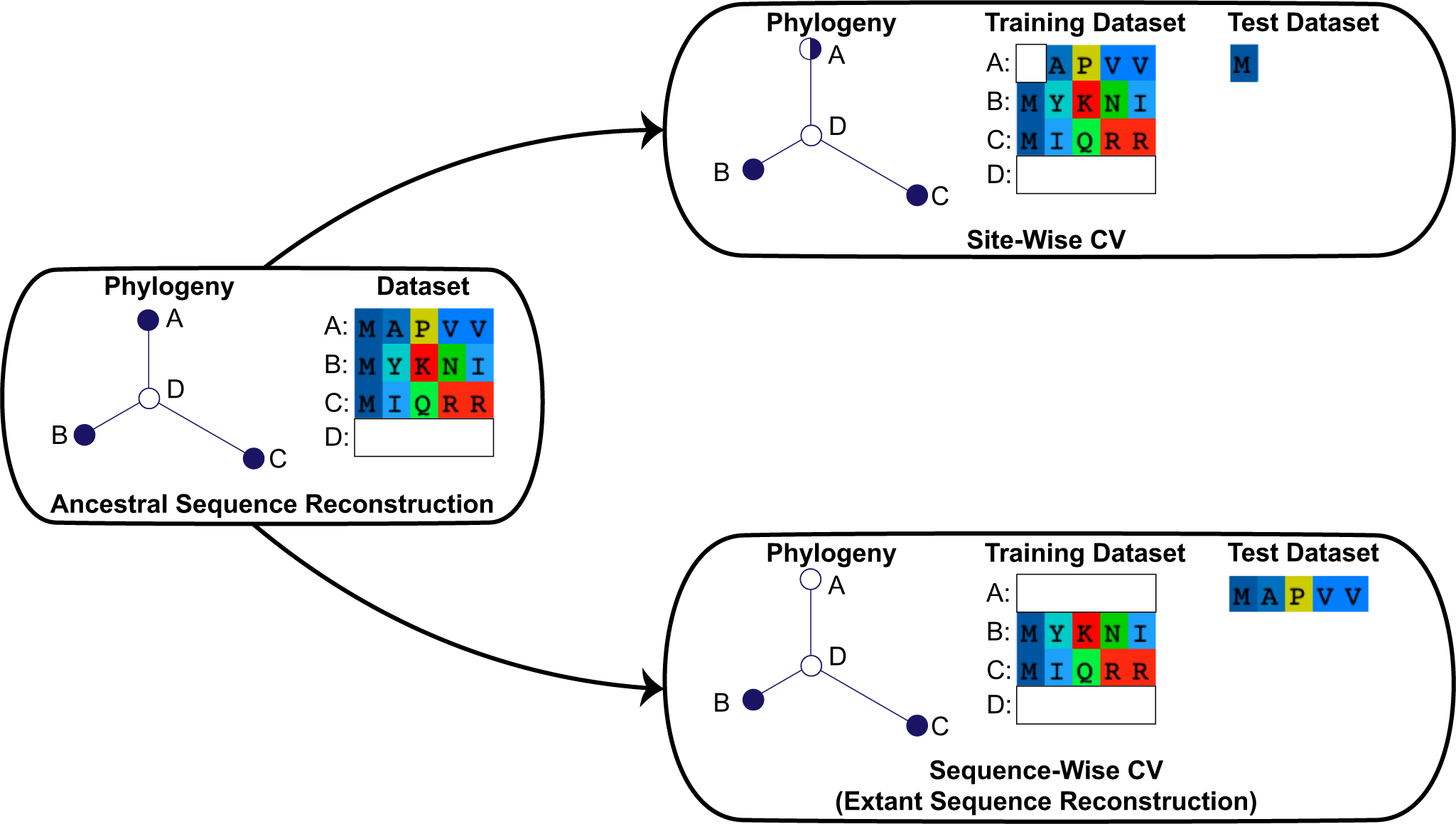
Cross-validation (CV) and Extant Sequence Reconstruction (ESR). On the left is a cartoon phylogenetic tree for a hypothetical alignment of three complete extant sequences (*A*, *B*, and *C*). The single hidden internal node has an unobserved ancestral sequence *D*. On the right are two types of CV explored in this paper. Each CV method has a phylogenetic tree inferred from the training dataset. The predictions of each CV method are benchmarked against the test set. Each tree node represents a sequence: filled circles represent complete observed sequences, empty circles represent unobserved sequences, and partially filled circles represent a site removed from that sequence.

This process is then repeated for every site in an extant sequence to generate a reconstructed distribution for the entire modern sequence *A*. The probability at every site in the sequence is calculated without knowledge of the true extant amino acid at that site. Thus, site-wise CV provides a probability distribution for an unobserved modern sequence conditional on the observed sequences in the alignment, just like conventional ASR.

One computational difference between ASR and site-wise CV is that ASR reconstructs every site using the same model parameters, whereas site-wise CV reconstructs each site using different model parameters. Because new ML model parameters are determined for each site, site-wise CV quickly becomes computationally intensive for common large biological datasets. For example, new ML model parameters must be inferred 19,377 times for site-wise CV of the kinase dataset, our smallest dataset, and the terpene synthase dataset is over an order of magnitude larger. Furthermore, to compare the reconstructions of different evolutionary models, this laborious process would need to be repeated for each model and for each protein family. To speed up the computation time, in all subsequent analyses we use a fast and accurate approximation to site-wise CV by employing a sequence-wise CV method that withholds one entire sequence at a time from the training set (see Methods, Supplementary Figure S8, and Fig. 3, bottom right). Hereafter we refer to such a reconstruction of a modern sequence found at a terminal phylogenetic node as ESR.

In the following we use ESR to evaluate reconstruction methodology when applied to the real biological sequence data in our three protein datasets. Qualitatively identical results were found when applying ESR to simulated data (Supplementary Figures S9-S11), but to economize in the main text we present and discuss the application and validation of ESR to experimental biological data, which is the primary advantage of ESR.

### Real biological dataset selection

We selected datasets from three protein families currently under investigation in our lab: (1) lactate and malate dehydrogenases (L/MDHs), (2) Abl/Src-related tyrosine kinases, and (3) terpene synthases (Table 1). These datasets were chosen because of their varying levels of taxonomic distribution and sequence divergence, and because they have unrelated topological folds presumably under different selection pressures. The L/MDH and Abl/Src datasets each consist of enzymes with relatively similar functions and specificities, whereas terpene synthases catalyze a diverse array of chemical reactions and have high sequence diversity. Consequently, the terpene synthase alignment has a greater fraction of gaps in comparison to the L/MDH and Abl/Src alignments.

**Table 1:**
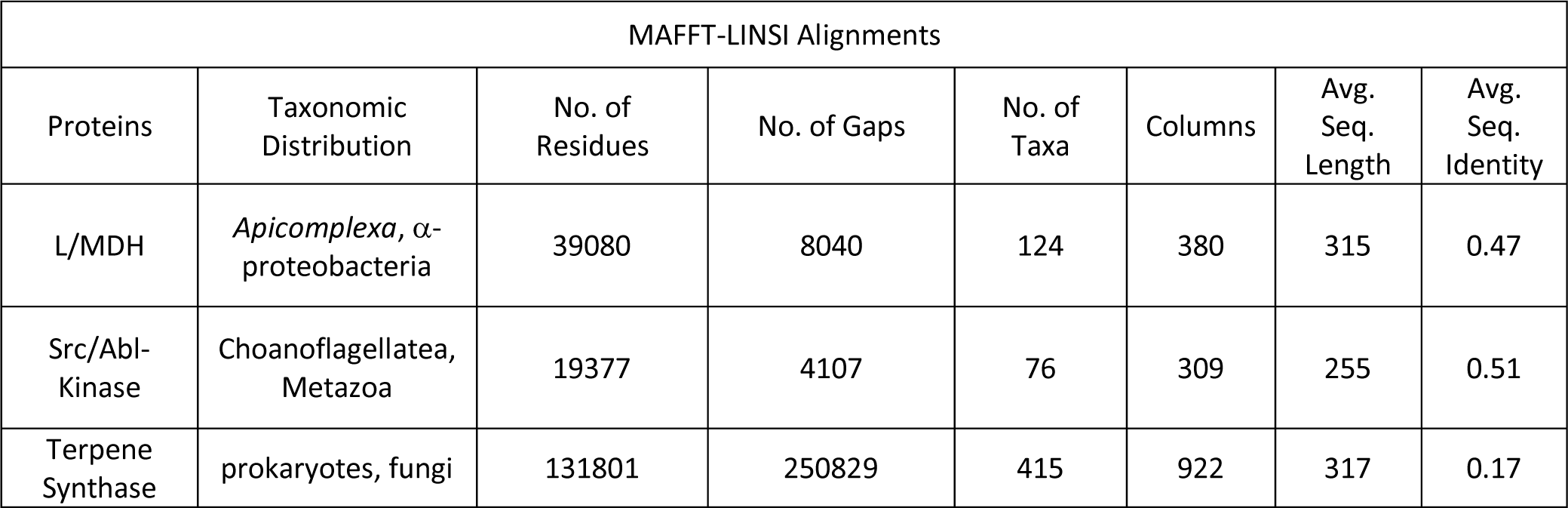
Summary of MAFFT-LINSI aligned datasets used in our CV analyses.

### Model selection: *LnL*, AIC, and BIC largely select similar models

ASR is fundamentally a problem of data prediction: based on the observed sequence data in an alignment, we wish to predict the true ancestral sequence for a given internal node in a phylogeny using a specific proposed model of sequence evolution. It would thus be useful to know which evolutionary model for our sequence data has the greatest predictive power. The predictive power of a model can be gauged by model selection criteria (Luo et al. 2010; Posada and Crandall 2001; Susko and Roger 2019), such as the Akaike Information Criterion (AIC) (eqn. 12) and the Bayesian (or Schwarz) Information Criterion (BIC) (eqn. 13) (Kalyaanamoorthy et al. 2017; Posada and Buckley 2004), which are commonly used in phylogenetics. AIC is a maximum likelihood (ML) method (Susko and Roger 2019) that aims to find the most predictive model, while the BIC aims to select the model that is most likely to be true given the observed sequence data (Neath and Cavanaugh 2012).

We calculated the AIC and BIC for our three protein datasets to evaluate the predictive performance of various competing models of evolution with increasing complexity (Table 2). As expected, more complex models resulted in higher raw maximum log-likelihood (*LnL*) scores, with the Poisson model having the lowest *LnL* and GTR20 with the highest. The AIC and BIC give scores that roughly track each other and the *LnL*. For both the L/MDH and kinase dataset, the AIC chooses the GTR+FO+G12 as the best evolutionary model, while the BIC chooses the LG+FO+G12 model (Table 2). However, in the terpene synthase dataset, both the AIC and BIC select GTR20+FO+G12, the most complex model. The differences between the AIC and BIC rankings are largely a philosophical matter; because we are most interested in sequence prediction, we hereafter refer to an evolutionary model preferred by the AIC criterion as a “better” or the “best” model.

**Table 2:** Summary of the model *LnL* and model selection criteria for various evolutionary models.

| L/MDH |  |  |  |  |  |  |
| --- | --- | --- | --- | --- | --- | --- |
| Model | Log-likelihood |  | AIC |  | BIC |  |
|  | Raw | From Best | Raw | From Best | Raw | From Best |
| Poisson+FQ | -30495 | -5227 | -30740 | -5018 | -31222 | -4951 |
| LG+FQ | -27087 | -1819 | -27332 | -1610 | -27815 | -1544 |
| LG+FO | -26628 | -1360 | -26892 | -1170 | -27412 | -1141 |
| LG+FO+G12 | -25484 | -216 | -25749. | -27 | **-26271** | **0** |
| LG+FO+G12+I | -25484 | -216 | -25750 | -28 | -26274 | -3 |
| GTR20+FO+G12 | **-25268** | **0** | **-25722** | **0** | -26616 | -345 |
| Abl/Src-Kinase |  |  |  |  |  |  |
| Model | Log-likelihood |  | AIC |  | BIC |  |
|  | Raw | From Best | Raw | From Best | Raw | From Best |
| Poisson+FQ | -23944 | -4176 | -24093 | -3967 | -24385 | -3897 |
| LG+FQ | -21674 | -1906 | -21823 | -1967 | -22115 | -1627 |
| LG+FO | -21277 | -1509 | -21445 | -1319 | -21774 | -1286 |
| LG+FO+G12 | -19988 | -220 | -20157 | -31 | **-20488** | **0** |
| LG+FO+G12+I | -19987 | -219 | -20157 | -31 | -20489 | -1 |
| GTR20+FO+G12 | **-19768** | **0** | **-20126** | **0** | -20827 | -399 |
| Terpene Synthase |  |  |  |  |  |  |

| Model | Log-likelihood |  | AIC |  | BIC |  |
| --- | --- | --- | --- | --- | --- | --- |
|  | Raw | From Best | Raw | From Best | Raw | From Best |
| Poisson+FQ | -198326 | -22010 | -199153 | -21801 | -201149 | -21297 |
| LG+FQ | -185083 | -8767 | -185910 | -8558 | -187906 | -8054 |
| LG+FO | -181614 | -5298 | -182460 | -5108 | -184502 | -4649 |
| LG+FO+G12 | -176978 | -662 | -177825 | -473 | -179869 | -17 |
| LG+FO+G12+I | -176971 | -655 | -177819 | -467 | -179865 | -13 |
| GTR20+FO+G12 | **-176316** | **0** | **-177352** | **0** | **-179852** | **0** |

### ESR SMP sequence probability accurately estimates the frequency of correct residues

To test the accuracy of our predicted reconstruction probabilities on actual biological data, we used ESR to reconstruct the extant SMP sequence at each terminal node of three phylogenetic trees and calculated the average probability of the SMP sequence. Because we know the true extant sequence at each terminal node, we compared the SMP reconstruction to the true sequence to ascertain the fraction of correctly predicted amino acids. As shown in Fig. 4a*-*c, the average probability of an extant reconstructed SMP sequence is an excellent estimate of the actual fraction of correct SMP residues. For each protein families and for every model of evolution, from least to most predictive, a plot of the average probability of the SMP reconstruction versus fraction correct yields a line with slope very close to unity. The average probabilities of the reconstructed extant sequences exhibit minimal systematic bias, regardless of the model, implying that extant reconstruction probabilities are robust to model misspecification. Similar results were found for ESR performed on simulated data (Supplementary Figures S9a, S10a, and S11a).

**Fig. 4:**
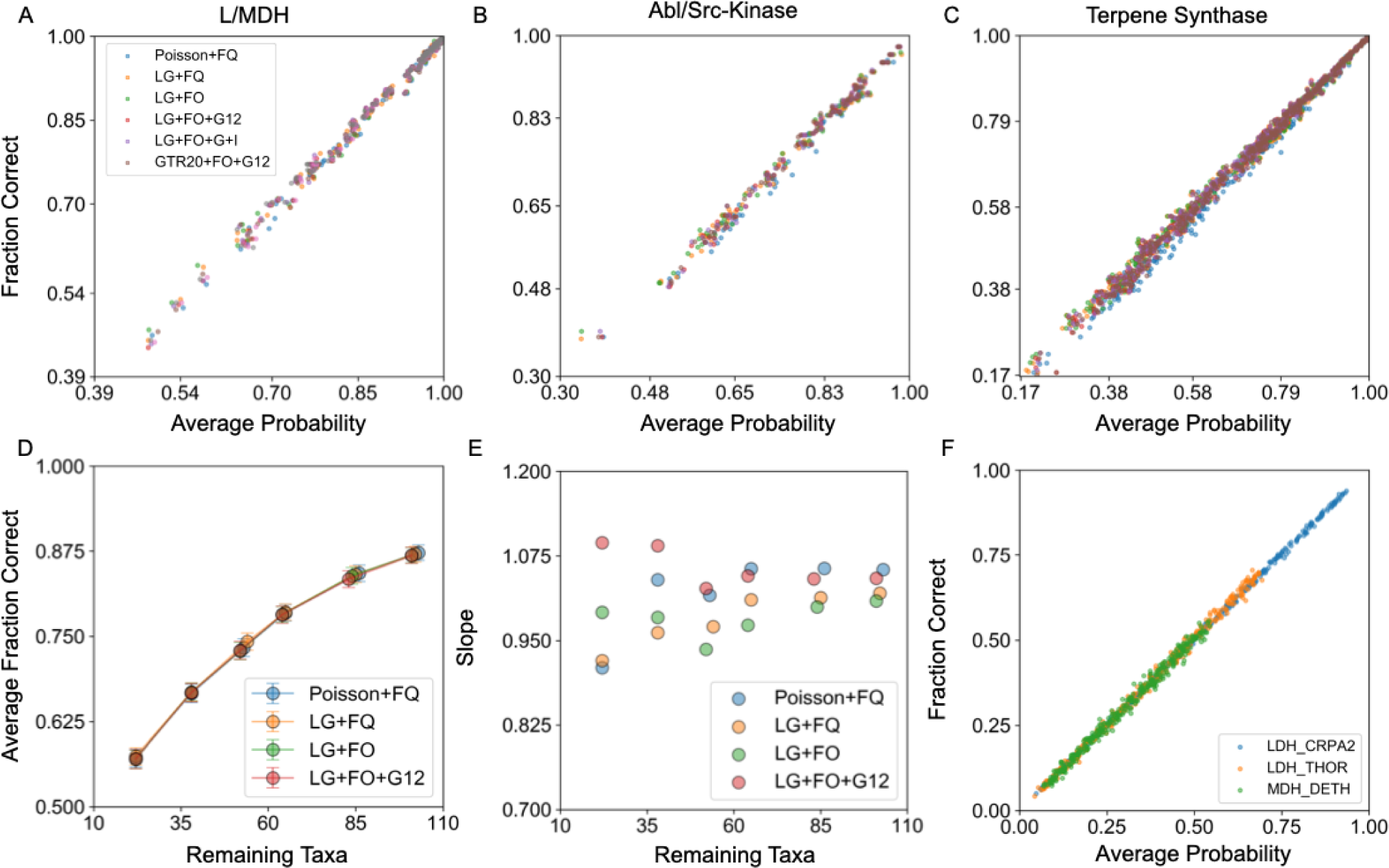
The average probability of extant reconstructions (ESR) accurately estimates the fraction of correct residues for real biological data. (*a*-*c*) Fraction correct vs. average probability of all SMP sequences for each model of evolution (L/MDH, kinase, and terpene synthase families). Each point represents a single SMP sequence from the reconstructed distribution. The slopes for linear fits of *a-c* are given in Supplementary Table 1. (*d*) Average fraction correct of all taxa vs. number of taxa in an L/MDH phylogeny, with taxa progressively removed and inferred using different models of evolution. (*e*) The corresponding slopes for linear regression of fraction correct vs expected fraction correct for each L/MDH phylogeny with different numbers of taxa removed. (*f*) Fraction correct vs. average probability of a set of sampled sequences for three different L/MDH proteins inferred using the LG+FO+G12 model of evolution. Each point represents a single sampled sequence from the reconstructed distribution.

### ESR residue probabilities retain accuracy for sparse taxon sampling, long branch lengths, and high uncertainties

Taxon sampling density is an important factor that influences the accuracy of phylogenetic analyses, including ASR (Heath et al. 2008; Salisbury and Kim 2001). Trees with sparse taxon sampling generally have longer branch lengths and more sequence diversity, which in turn decreases the certainty in ancestral state reconstructions due to less sequence information. Consequently, the high accuracy of ESR probabilities could degrade with more diverse datasets.

To assess the effect of taxon sampling on the accuracy of ESR probabilities, we repeated ESR on our L/MDH dataset after progressively pruning terminal nodes with short branch lengths, resulting in trees with increasingly longer branch lengths. The average fraction correct for SMP reconstructions decreases as sequences are pruned from the tree, indicating the reconstructions get more uncertain as expected (Fig. 4d). However, the linear relationship between fraction correct and average posterior probability holds despite the longer branch lengths in these pruned datasets (Fig. 4e). Similarly, the terpene synthase dataset includes many SMP sequences below 50% average probability, which accurately predict the fraction correct as low as 17% (Fig. 4c).

SMP residue probabilities, as extreme values, may be somehow unusually accurate, and lower, non-SMP residue probabilities may be less accurate. To address this concern, we chose three enzymes from our L/MDH dataset and sampled sequences from their respective reconstructed distributions. We chose LDH_CRPA2, LDH_THOR, and MDH_DETH because they spanned a wide range of average probabilities for their respective SMP sequences. The average probabilities of the SMP sequences are 95%, 80%, and 65% respectively. We used biased sampling from the ancestral distributions to generate increasingly lower average probability sequences. Like with the SMP sequences, a plot of the fraction correct in a sampled sequence against its average probability gives a line of slope close to one (Fig. 4f). ESR probabilities from low average probability sequences (0.05) to very high average probability sequences (0.95) accurately estimate the frequency correct. Overall, the probabilities of extant reconstructions are just as accurately calculated regardless of the level of uncertainty.

### For ESR, better models also can decrease SMP probabilities and fraction correct

Like the results from ASR with simulated data, for ESR the total *LnP* of SMP sequences is anticorrelated with the true sequence total *LnP* and model complexity for all three protein families, in line with our observations from simulated data (Fig. 5a and 5b, Supplementary Figures S9c, d, S10c, d, and S11c, d). Similarly, the model *LnL* is anticorrelated with SMP sequence *LnP* (Table 2, Supplementary Table 6).

**Fig. 5:**
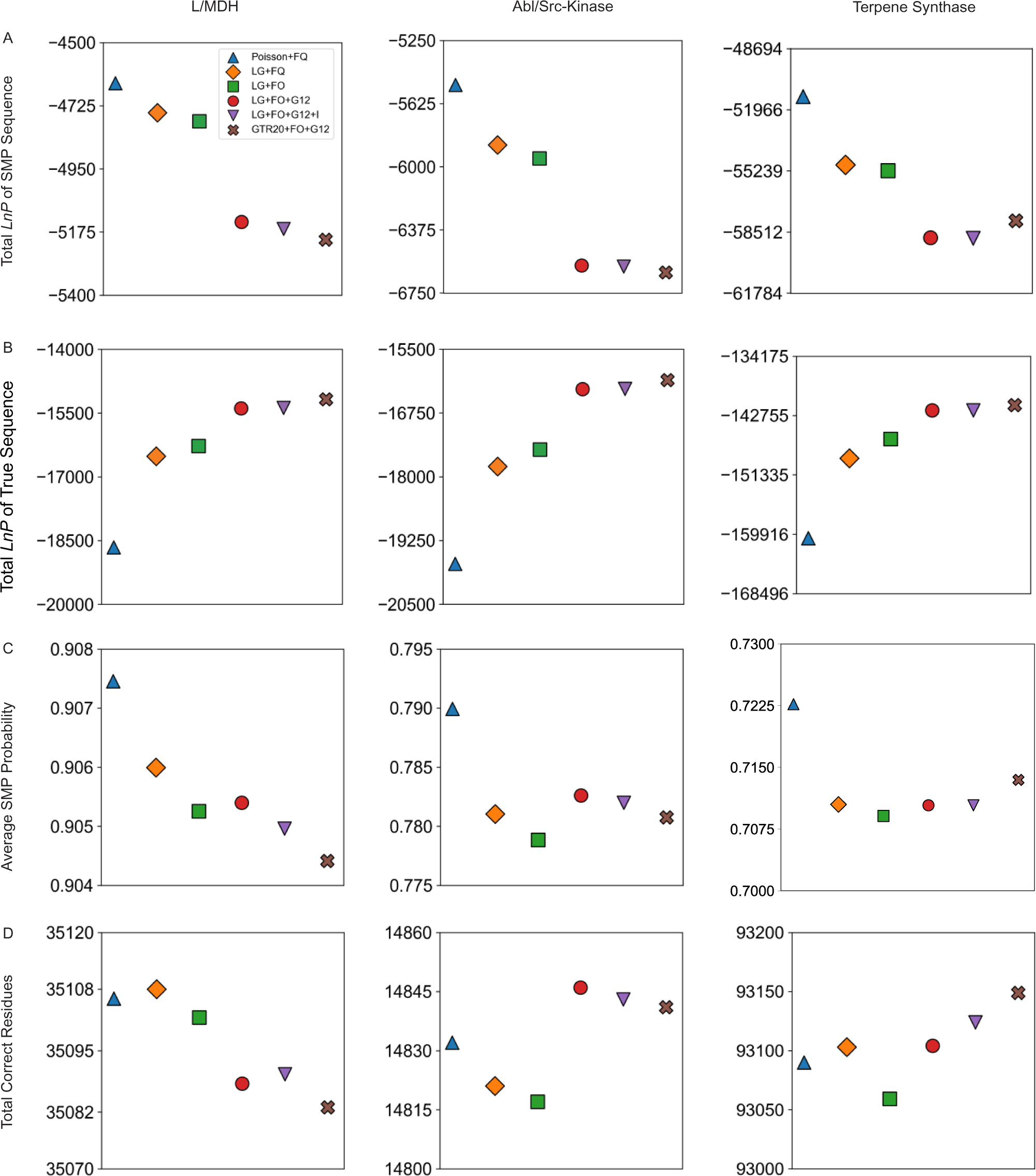
Extant SMP probabilities and number of correct residues generally decrease with better models. (*a*) Total *LnP* of extant SMP sequences in a phylogeny for each model of evolution. (*b*) Total *LnP* of true extant sequences for each protein family and model of evolution. The values for (a) and (*b*) are given in Supplementary Table 7. (*c*) The average of the average posterior probability for all reconstructed SMP sequences and (*d*) the total number of correct amino acids in the reconstructed SMP sequences for each model.

Similarly, the average SMP sequence probability generally decreases as model complexity and predictiveness increases (Fig. 5c). A lower average probability should translate to fewer correct amino acids in the SMP reconstruction. However, the differences among the models in the number of correct residues is relatively modest in comparison to the total number of correct residues (Fig. 5d). For perspective, the L/MDH dataset contains over 35,000 residues in total, but only 18 more residues are correctly predicted by the Poisson+FQ model compared to the LG+FO+G12 model (amounting to one additional correct residue per seven sequences on average).

We note that the effect of adding additional explanatory model parameters is not completely consistent across the protein families. For instance, adding among-site rate variation (+G12) to the evolutionary model increases the number of correct amino acids for only the kinase family. Nevertheless, the SMP reconstructions derived from worse evolutionary models, as judged by predictive model selection, tend to have higher average probabilities and more correct amino acids than reconstructions from better models.

### For ESR, better models also improve true residue probabilities at a cost to SMP residues

Both the ASR and ESR results suggest that there is a tradeoff between true and SMP residue probabilities. To test this, we plotted a histogram of the true residue probabilities at sites in which the SMP residue is incorrect for the L/MDH protein family (Fig. 6a). True residues at incorrect sites are skewed toward low probabilities regardless of evolutionary model for our L/MDH dataset, yet the probabilities of the true residues clearly increase for more predictive models. From the histogram of SMP residue probabilities at incorrect sites, we see that both the mean and skew decrease as the complexity of the evolutionary model increases for all three protein families (Fig. 6b and Supplementary Figures S12-S13). Hence, in general, more predictive models increase the probability of the true residue at the expense of the SMP residue.

**Fig. 6:**
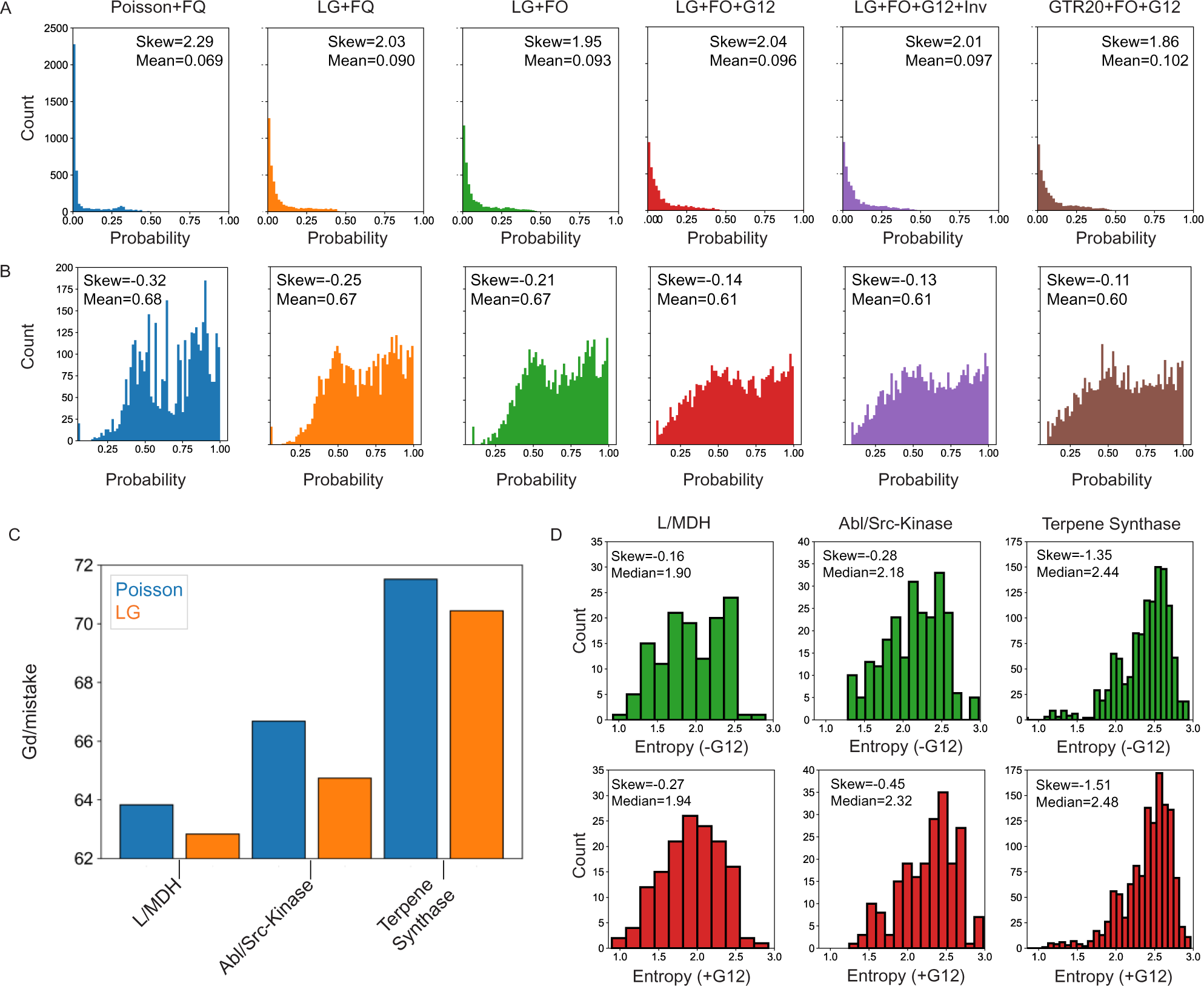
Better models make better mistakes. (*a*) Histograms of the true residue probability at incorrect sites for each model of evolution for the L/MDH dataset. (*b*) Histograms of the SMP residue probability at incorrect sites. (*c*) The total Grantham distance per mistake for Poisson+FQ and LG+FQ models. (*d*) Histograms of column entropy for each unique mistake in the LG+FO (upper, green) and LG+FO+G12 (lower, red) models for the three protein families. The skew and median of each histogram is shown.

### Better substitution matrices produce SMP sequences that are more chemically similar to the true sequences

Why should the total number of correct residues decrease despite improvement in the model *LnL* or AIC? While the absolute number of correct amino acid residues is a common and simple measure of the distance between two sequences, not all amino acid mistakes in a sequence are biologically equivalent. Some mistakes are more detrimental to protein function than others, due to differences in the chemical properties of amino acids or the strength of functional constraint at a given site. A reconstructed sequence with many benign mistakes nevertheless may be chemically and biophysically closer (and hence functionally more similar) to the true sequence than a sequence with only a few highly perturbing mistakes.

An evolutionary model incorporating a substitution matrix with unequal amino acid exchange rates (like LG) can account for biophysical dissimilarities in amino acid substitutions, and therefore such a model should make fewer mistakes of high chemical detriment yet allow more mistakes that are chemically similar. To test this, for the Poisson+FQ and LG+FQ evolutionary models we calculated the chemical dissimilarity between reconstructed and true sequences using the Grantham distance (*Gd*). The *Gd* is a common pairwise metric for amino acid biophysical dissimilarity that is a function of volume, hydrophobicity, and number of heteroatoms. (Grantham 1974) The *Gd* per mistake in the SMP sequence indeed decreases substantially for all models that incorporate the LG substitution matrix in place of the Poisson equal rates assumption (Fig. 6c). Thus, better evolutionary models with realistic amino acid substitution processes may produce reconstructed SMP sequences that have lower average probability and make more naïve mistakes, but overall these mistakes are more chemically conservative.

### Rate variation reduces the number of mistakes at conserved sites

Evolutionary models with rate-variation across sites can account for differences in functional constraint. A site that is functionally important generally has a lower rate of substitution and divergence compared to a site that has low functional constraint. When reconstructing a sequence, a mistake at a slow evolving site is more likely to be deleterious than a mistake at a fast-evolving site. We thus expect that including among-site rate-variation in an evolutionary model may increase the number of mistakes at sites with high rates while reducing the number of mistakes at sites with low rates.

To test this hypothesis, we quantified the divergence at sites with mistakes in the SMP sequence using the amino acid entropy, a measure of site conservation. In general, highly variable, fast-evolving sites have a high entropy, while conserved, slow-evolving sites have a lower entropy. We compared the LG+FO and LG+FO+G12 models by plotting the distribution of the entropies at sites with mistakes (Fig. 6d). For each protein family the median and skewness of the entropy distribution increases when incorporating rate variation among sites, indicating that models which include rate-variation make fewer mistakes at functionally constrained, conserved sites while making more mistakes at less functionally important sites. Hence, for SMP sequences, models with rate variation may make more mistakes in general, but the mistakes likely have less of a functional impact overall.

### Increasing model predictiveness improves the expected log-probability and the expected chemical similarity to the true sequence

In addition to the SMP-based statistics, we also asked how expected values behave as a function of the evolutionary model. Unlike the SMP *LnP*, the *eLnP* generally improves with model predictiveness and tracks with AIC (Fig. 7a). This result is consistent with our results for the *eLnP* of reconstructed extant sequences with simulated data (Supplementary Figures S9e, S10e, and S11e). Additionally, the expected number of correct sites improves for all three protein families when among-site rate variation is included in the models (Fig. 7b). As with the *Gd* for the SMP sequences, we see that changing the substitution matrix from Poisson to LG greatly improves the expected *Gd* (Fig. 7c). The expected *Gd* also tends to monotonically decrease with increasing model predictiveness. Hence, both ASR and ESR results suggest that for evaluating ancestral reconstructions expected value statistics are more useful than statistics based on the SMP.

**Fig. 7:**
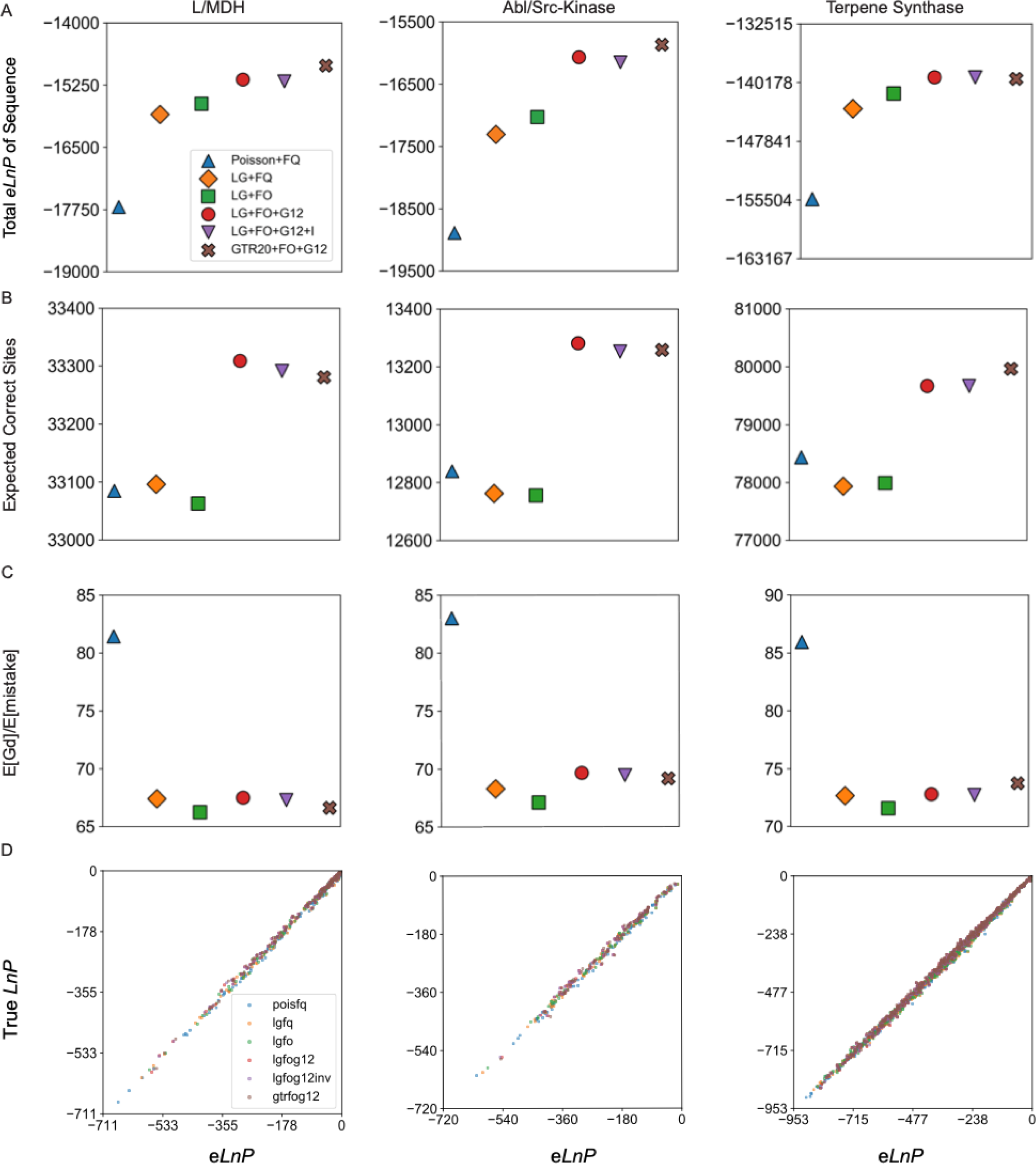
The expected *Gd* for ESR is generally improved by increasing model complexity. (*a*) Total *eLnP* of the extant sequence for each model of evolution and each protein family. The values for (*a*) are given in Supplementary Table 6. (*b*) The expected number of correct amino acids of the reconstructed distribution and (*c*) the expected Grantham distance per expected number of mistakes for different evolutionary models. (*d*) The true sequence *LnP* plotted vs. *eLnP* for each protein family and model of evolution. Each dot represents a reconstructed SMP sequence. Linear regression slopes for (*a*) are given in Supplementary Table 1.

### The expected *LnP* is an accurate estimate of the true sequence *LnP* for extant reconstructions

We expect that if our evolutionary models are capturing important features of real evolutionary processes, then the extant *eLnP* should accurately approximate the *LnP* of the true extant sequence. We find that the extant *eLnP* for a given sequence is an excellent estimate for the true extant sequence *LnP* for both real biological data and simulated data regardless of evolutionary model (Fig. 7d, Supplementary Figures S9f, S10f, and S11f). Like the fraction correct vs. average probability, linear least-squares fits of the true sequence *LnP* vs. *eLnP* gives a slope of approximately 1.0 for all models of evolution and each protein family (Supplementary Table 1). Hence, we can accurately estimate the *LnP* of the true extant sequence at a hidden node from the reconstructed distribution using the e*LnP*, without knowing the true sequence.

## Discussion

### More accurate models give more accurate ancestral probabilities

Several recent studies have suggested that phylogenetic model selection does not matter as the resulting inferences are all equally distant from truth (Abadi et al. 2019; Spielman 2020; Tao et al. 2020; Williams et al. 2006). Our analyses of simulated data, however, corroborate previous studies indicating that accurate ancestral sequence reconstructions require accurate models (Del Amparo and Arenas 2022; Finnigan et al. 2012; Hanson-Smith et al. 2010; Zhang and Nei 1997). When the model is misspecified, ancestral SMP probabilities are inaccurate and biased, yet the probabilities become increasingly less biased and more accurate as the model approaches the true generating model (Figures 2, 3, and Supplementary Figures S1, S2, S4, S6 and S7).

Why does GTR20+FO+G12 provide accurate probabilities for all simulations despite being overparameterized? The GTR20+FO+G12 model is correctly specified for the data, since both the Poisson+FQ and LG+FO+G12 models are nested within the GTR20+FO+G12 model. The fitted parameters of the GTR20+FO+G12 model will approximate those of the Poisson+FQ and LG+FO+G12 with higher certainty as the data set size increases. In contrast, the overparameterized LG+FO+G12 model is misspecified for datasets simulated under the Poisson+FQ model and cannot converge to the true model regardless of the amount of data. While using an overparameterized model tends to be less problematic than using an underparameterized model, especially with large datasets, not all overparameterized models are equally useful, and model selection is necessary to pinpoint the best model for ASR.

### Extant reconstruction probabilities are accurate even for misspecified models

Before applying ESR to real proteins, we compared the behavior of ESR and ASR using simulated data. We performed ESR 3075 times by fitting 5 different models on three different datasets simulated using the LG+FO+G12 model of evolution. For both ESR and ASR, the probability of SMP residues generally decreases as the fitted model becomes more similar to the true LG+FO+G12 model (compare figs. 1b-c, S1b*-*c, and S2b*-*c with S9b*-*c, S10b*-*c, and S11b*-*c, respectively). Likewise, for both extant and ancestral reconstructions, the SMP sequence *LnP* and the true sequence *LnP* are anticorrelated for under-parameterized models (compare figs. 1c-d, S1c-d, and S2c-d with S9c-d, S10c-d, and S11c-d, respectively). This behavior is expected since ESR is simply ASR methodology applied to terminal nodes rather than the more conventional internal nodes.

The absolute accuracy of reconstruction probabilities is, however, one notable difference between ESR and ASR. The average probability of an SMP sequence from ASR of simulated data is only an accurate estimate of fraction correct with a correctly specified model (Fig. 1a and Fig. 4a-c). A misspecified model results in ancestral reconstructions that overestimate their expected fraction correct. Unlike ASR, with ESR the calculated average probability of the SMP sequence is a reliable estimate of true fraction correct regardless of evolutionary model (Supplementary Figures S9-S11). Similarly, with ESR the calculated *eLnP* is a reliable estimate of the true sequence *LnP* regardless of model. Nevertheless, the ESR probabilities are also slightly biased, just less biased than ASR probabilities. This is shown by the regression of actual fraction correct versus average sequence probability for misspecified models, which deviate from a slope of 1.0 and y-intercept of 0 significantly more than correctly for specified models (Supplementary tables 1 and 6).

Notably, for all ancestral sequence reconstructions, fraction correct appears to be a monotonically increasing function of the average probability of the SMP sequence, indicating that even inaccurate reconstructed probabilities are useful as a gauge of relative accuracy. Furthermore, the reconstructed probabilities become more accurate and unbiased as they increase and approach a probability of 1.0 (Figures 1, 2, and Supplementary Figures S1, S2, S4, S6, and S7). For reconstructions with average probability, say, ≥ 0.95, the inaccuracies in the probabilities are minimal for even misspecified models. As more evolutionarily relevant model parameters are included, the expected fraction correct approaches the actual fraction correct, which underlines the importance of model selection criteria to reduce bias in ancestral probabilities.

### Sequence correlations compound bias in ancestral reconstructions, but not in extant reconstructions

Why are ESR probabilities more accurate than ASR probabilities for misspecified models? The probabilities for extant sequences determined by ESR are calculated using the same theoretical methodology as ancestral probabilities in ASR. However, the terminal nodes used in ESR differ from the internal nodes that are the focus of conventional ASR. Internal nodes are connected to and receive information from three other nodes, whereas a terminal node is connected directly to only a single internal node. All else equal, the probabilities at a terminal node are calculated using less information than an ancestral node. Given these differences we expected that the behavior of reconstructed extant sequences might differ from reconstructed ancestral sequences.

We see that overall the extant reconstructions are much lower in average sequence probability than the ancestral reconstructions. For example, the average SMP ancestral sequence probability is approximately 0.9 for the Poisson+FQ reconstructions in the simulated terpene synthase dataset, whereas for the extant reconstructions the average SMP sequence probability is 0.68. This is consistent with the fact that ancestral sequences receive more information from three branches, whereas tips receive less information from a single branch.

Phylogenetic structure quantifies correlations between residues in different sequences at a site in an alignment. Different sequences are not independent data, and the correlations are modelled by the phylogenetic tree. Internal nodes receive information on sequence correlations from three different domains of a tree, while terminal nodes only receive information through one branch. Since a terminal node has only one source of information, there is no potential for conflicting information coming from multiple sources. Incorrectly modeled correlations, for instance, from a misspecified model, should therefore affect and bias reconstructions at internal nodes more than terminal nodes, and this is apparently reflected in the more biased ASR probabilities compared to ESR probabilities.

### Better models result in SMP sequences more chemically similar to truth

How can the true model result in a less probable SMP sequence that is expected to result in fewer correct residues? A general answer is that ML methodology does not maximize the number of correct residues; rather, ML maximizes the probability of the observed sequence data by finding the best value of each model parameter. Similarly, model selection methods like AIC maximize the expected probability of the observed sequence data by finding the best model. In light of this, we should expect that using ML with a more predictive model may result in lower probability SMP sequences with fewer total correct amino acids. Including parameters that increase the AIC improves the predictiveness of the model by capturing an influential aspect of the evolutionary process. Biologically meaningful model parameters produce predictions that are more biologically and evolutionarily realistic, but more biologically realistic sequences may not necessarily have fewer amino acid errors.

For instance, the evolutionary impacts of amino acid substitutions are not all equal because of differing degrees of chemical similarity among amino acids (Norn et al. 2021). Therefore, replacing a Poisson substitution matrix with the LG substitution matrix should have an impact on SMP biophysics by skewing towards mistakes that are chemically similar to the true amino acids. These considerations suggest that the absolute number of correct residues is the wrong metric to assess how substitution matrices impact SMP reconstructions. Using the Grantham distance *Gd* as a metric to judge the difference between SMP reconstructions and the truth, we saw that the LG substitution matrix improves the types of mistakes made in the SMP sequence (Fig. 6c).

Similar to a substitution matrix, among-site rate variation in a phylogenetic model phenomenologically accounts for differences in purifying and adaptive selection among sites in a protein. Sites with higher functional constraint generally evolve with slower rates and result in alignment columns with lower amino acid entropy. Hence, a mistake at a low entropy site is not on average functionally equivalent to a mistake at a high entropy site; rather, a mistake at a low entropy, functionally constrained site is more likely to be functionally detrimental. We see this effect in the mean and skew of the entropy histogram of unique reconstruction mistakes when we compare models with and without rate variation (Fig. 6d). The differences between models may appear to be small in terms of *Gd* per mistake and the change in mean entropy, yet the benefits of making better mistakes can be critically important. Even a single chemically detrimental amino acid change, or a single conservative change in an active site, can have catastrophic effects on protein function.

These examples highlight what is likely a general consideration in evaluating the effects of model selection methodologies: when evaluating the “closeness” of model inferences to truth, it is critical to choose the proper metric. For our phylogenetic models, the absolute number of differences between a reconstructed sequence and the true sequence is an overly simplistic metric for quantifying biophysical accuracy. Similarly, when evaluating the closeness of an inferred phylogeny to the true phylogeny, the raw Robinson–Foulds distance is likely the wrong metric for quantifying evolutionary accuracy (Abadi et al. 2019; Spielman 2020; Tao et al. 2020; Williams et al. 2006). Rather, more evolutionarily meaningful metrics like a generalized Robinson–Foulds distance may be more appropriate (Smith 2021).

### Better models improve expected properties of the reconstructed distribution

The SMP sequence is typically resurrected and used to provide conclusions about protein evolution, primarily because of its experimental tractability (Akanuma et al. 2013; Boucher et al. 2014; Chang et al. 2002; Gaucher et al. 2003; Nguyen et al. 2017; Thornton et al. 2003; Wilson et al. 2015). However, the SMP sequence is known to be an unusual and biased sequence, and as a single point estimate of the true ancestral sequence its use discards information in the ancestral distribution on the uncertainty in reconstructed amino acid states (Krishnan et al. 2004; Matsumoto et al. 2015; Williams et al. 2006; Yang 2006). A more theoretically justified method is to study the expected properties of the proteins described by the reconstructed distribution. In some cases the expectations can be calculated analytically (as is the case with all examples in this work), but when that is infeasible the expectations can be approximated experimentally by sampling sequences from the ancestral distribution and studying the average properties of the samples (Gaucher et al. 2008). For these reasons we wished to explore the effects of model parameters on the expected properties of the entire reconstructed distribution.

Changing the substitution matrix from Poisson to LG consistently lowers the expected *Gd* per expected mistake for all three protein families without adding additional parameters to the model (Fig. 7c). The LG substitution matrix improves the probability of true residues at incorrect sites of the SMP sequence (Fig. 5c), while decreasing the probability of the SMP residue at incorrect sites (seen from the slight change in histogram skew of the probabilities of incorrect amino acids) (Fig. 5d). Even though changing the substitution matrix improves the chance of selecting the true residue, it has a lesser effect on reducing the chance of the wrong SMP residue. This could explain why the expected *Gd* per mistake improves significantly, but the expected number of correct residues does not improve.

Adding among-site rate variation to the model results in the largest and most consistent improvement to the expected number of correct residues (Fig. 7b). Among-site rate variation is the biggest factor in reducing the probability of incorrectly guessed SMP residues (Fig. 6b), while there is not as much improvement in the probability of the true residue (Fig. 6a).

When resurrecting an ancestral protein, we are ultimately most interested in accurately reproducing the biophysical and functional properties of the true ancestor, rather than simply getting the sequence correct. The *Gd* and the total number of correct residues are proxies for the experimental properties of resurrected enzymes. Having more correct residues and a lower *Gd* are both likely to be result in a reconstructed protein with biophysical properties that are closer to the true ancestor. However, the ultimate judge is experimental comparison of reconstructed and true biophysical properties, which we leave to future work.

### SMP reconstructions have the highest expected identity to the true sequence

One justification for using the SMP reconstruction is that it is expected to have the fewest mistakes, relative to the true ancestor, of all possible sequences. We have found that the frequency of successfully predicting a correct amino acid is positively correlated to that amino acid’s reconstruction probability. Hence using a lower probability sequence should generally result in fewer correct amino acids predicted than the SMP sequence. We indeed find that fewer correct residues are expected in randomly sampled sequences (Fig. 7b) relative to the SMP sequence, for all protein families across all models of evolution (Fig. 6b). In addition, sampled sequences are expected to be less chemically similar to the true sequence (Fig. 7c) than the SMP sequence is to the true sequence (Fig. 6c). Taken together, the SMP reconstructions result in fewer errors and more chemically similar sequences on average than sequences sampled from the posterior probability distribution.

### Biased sampling of reconstructions may miss many true residues

The SMP sequence is just one possible sequence out of the reconstructed distribution. The reconstructed distribution contains information for constructing all other possible sequences of the same length, with some being more plausible than others. There are three widely used methods for generating plausible alternatives to the SMP sequence: (1) unbiased random sampling of a residue at each site, (2) biased random sampling of a residue at each site, and (3) deterministic generation of lower probability sequences, such as the “AltAll” method (*e.g.*, select the second most probable residue at a site when its probability is greater than 0.2) (Akanuma et al. 2013; Gaucher et al. 2008; Lim et al. 2016; Wheeler et al. 2016). The latter two methods essentially assume that reconstruction distributions do not contain correct yet low probability (e.g., < 0.2) residues.

To determine if AltAll and biased sampling miss low probability yet correct residues, we investigated the distribution of true residue probabilities at incorrect SMP sites. We find that, when the SMP chooses the wrong residue, the probability of the true residue at that site is highly skewed towards 0.05 (Supplementary Figure S10). Therefore, restricting residue sampling to a probability >0.2 will miss many true residues, because biased sampling generates incorrect residues at a frequency higher than unbiased sampling.

### Sampled reconstructions can be more accurate than the SMP reconstruction

Of all possible sequences, the SMP sequence has the fewest expected number of differences from the true sequence. Unbiased sampling from the reconstructed distribution generates sequences that have more expected mistakes than the SMP sequence. Previously, Eick *et al*. visually represented sampled sequences by plotting them as the difference in *LnP* vs the number of amino acid differences, both relative to the SMP reconstruction (Eick et al. 2017). The main purpose of these plots was to illustrate that sampled sequences may be non-functional because of their lower probability and greater number of expected errors relative to the SMP. (Note that our terminology follows that of (Matsumoto et al. 2015)). Eick *et al*. call the “ML reconstruction” what we call the SMP reconstruction, and they call “Bayesian sampling” what we refer to as sampling from the ML reconstruction distribution.)

Using ESR, we reconstructed the SMP sequence and 10,000 sampled sequences for three extant proteins (LDH_CRPA2, LDH_THOR, and MDH_DETH) that represent a wide range of average posterior probabilities for their respective SMP sequences (0.95, 0.80, and 0.65 respectively). Like Eick et al, we plotted the *LnP* of each reconstructed sequence against the number of differences relative to the SMP sequence (Fig. 8a*-*c). As previously seen, the sampled sequences all have lower *LnP* than the SMP sequence (which is true by definition of the SMP). Furthermore, the sampled sequences form a cloud with varying numbers of sequence differences from the SMP, and it is clear that the SMP is an extreme outlier from the reconstructed distribution.

**Fig. 8:**
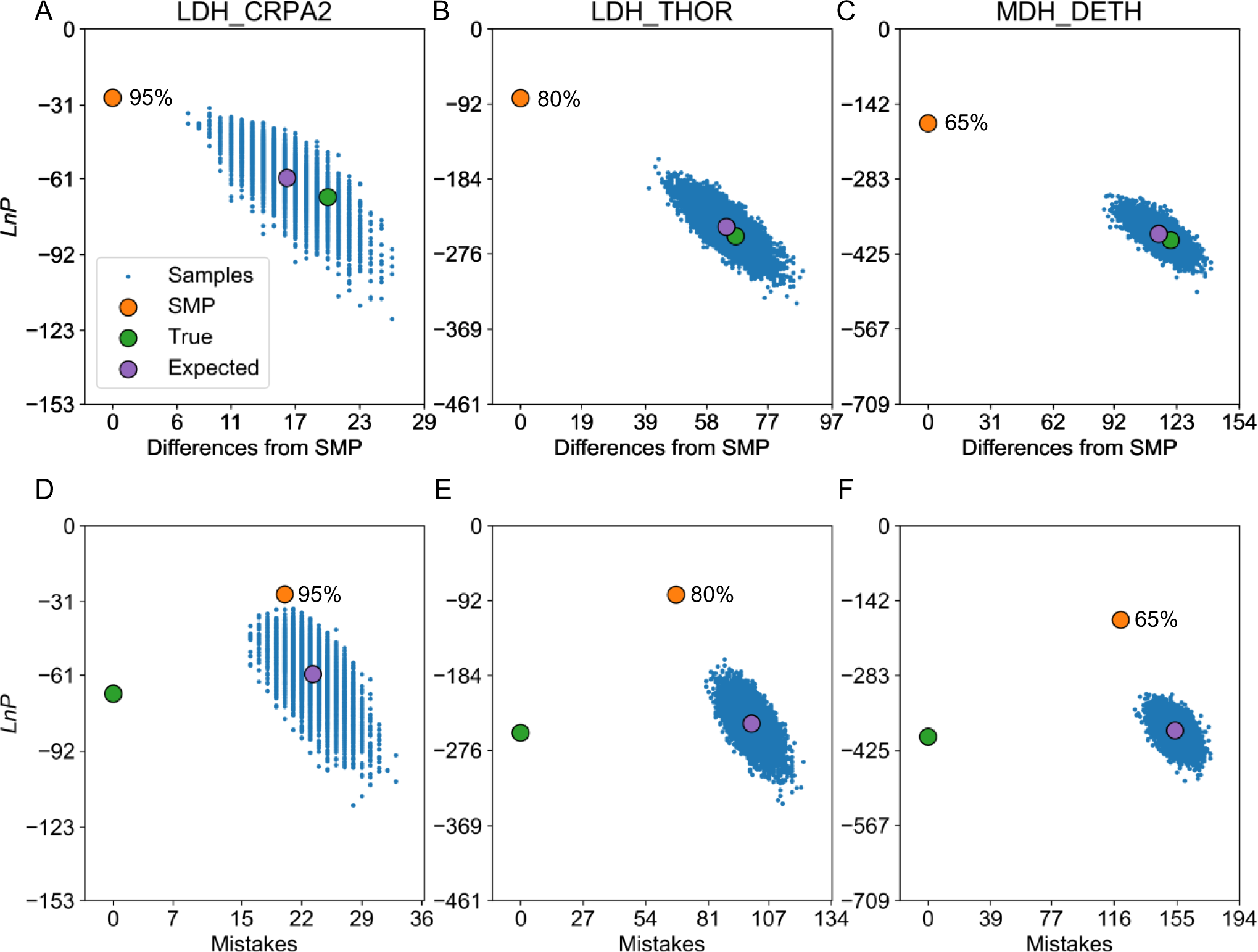
The SMP sequence may have significantly more mistakes than sampled sequences. Sampled sequences (blue dots) and other specific sequences (colored circles) for an ESR analysis of three different extant L/MDH proteins. The blue dots represent 10,000 sequences sampled from the reconstructed distribution, the green circle is the true sequence, the orange circle is the SMP sequence, and the purple circle is the expected property of the reconstructed distribution. (*a-c*) The *LnP* of a sequence vs the number of differences from the SMP for three different proteins. (*d-f*) The *LnP* of a sequence vs the number of mistakes in a sequence (relative to the true sequence) for the three different proteins. The SMP sequence average probability is shown next to the SMP sequence (orange dot).

Because our data are from an ESR analysis, we know the identity of the true sequence, and so we are also able to plot values for the true sequences (green dots) and the expected values (purple dots) from the reconstruction distribution. It is immediately apparent that the true sequence has *LnP* typical of the sampled distribution, unlike the SMP reconstruction. The expected *LnP* of the reconstructed distribution (purple dots, y-axis) is very close to the *LnP* of the true sequence (green dots, y-axis), indicating that we can accurately approximate the true sequence *LnP* without knowing the identity of the true sequence. Note that for sampled sequences the expected number of differences from the SMP sequence is equal to the expected number of mistakes in the SMP sequence (purple dots, x-axis in figs 8a*-*c), which can be calculated from the average probability per residue of the SMP sequence (Eqn 8 in Methods). Because we know the true sequence, we can also plot the actual number of mistakes in the SMP sequence for comparison (green dots, x-axis in figs 8a*-*c). As can be seen in the plots, the expected number of mistakes in the SMP is strikingly close to the actual number of mistakes in the SMP, as anticipated.

These plots (Fig. 8a-c) could give the impression that the sampled sequences have many more mistakes than the SMP. However, the number of amino acid differences from the SMP is not the same as the number of mistakes, which are differences from the true sequence. In principle a sampled sequence can be very close in identity to the true sequence yet differ greatly from the SMP. Thus, it would be more informative to use the true sequence as the benchmark for comparison, by plotting differences from the true sequence rather than from the SMP sequence. Though this is impossible with a standard ASR analysis, it is possible with ESR.

In figs 8d*-*f, we plot the same ESR data but as differences from the true sequence. As before, the sampled sequences all form a cloud of similar *LnP* and mistakes, and the expected values sit in the middle of this cloud. The expected number of mistakes in a sampled sequence (purple dot) is the expected average probability of the distribution of reconstructions (calculated from Eqn 9 in Methods). In these plots, the SMP sequence is much closer to the cloud of sampled sequences, and the difference in number of mistakes between the sampled sequences and the SMP is much lower than the difference between the sampled sequences and the SMP as shown in the Eick-style plots.

For LDH_CRPA2, many sampled sequences (∼11%) in fact have fewer mistakes than the SMP sequence (e.g., all the blue sampled sequences shown to the left of the orange SMP, which has 20 mistakes). The number of mistakes in sampled sequences follows a Poisson-Binomial distribution roughly centered on the expected number of mistakes (for LDH_CRPA2 mean = 23, sd = 2.3). In general, many sampled sequences may have fewer mistakes than the SMP sequence when the SMP has high average probability (e.g., greater than ∼0.9). We do not see this for LDH_THOR and MDH_DETH because the number of mistakes in their SMP sequences is far fewer than the expected number of mistakes for the distribution.

### ESR highlights relative information content for regions in a phylogeny

The many different methods for calculating branch supports provide a measure of confidence in the existence of a specific branch, and by extension, a specific ancestral node (Anisimova and Gascuel 2006; Anisimova et al. 2011; Minh et al. 2013). However, a branch support does not provide information about the quality of a sequence reconstruction at that node. The *eLnP* of a sequence becomes an increasingly accurate estimate of the true sequence *LnP* as the model becomes more correctly specified. The *eLnP* of all internal and external nodes can be included in a phylogeny, as a complement to branch supports, to indicate and visualize which regions have high information and which have low information about ancestral and extant nodes. As an example, for an Apicomplexa L/MDH dataset we calculated the normalized *eLnP* for all nodes and mapped them onto the phylogeny (Fig. 9). We can see that there is a specific clade, composed of bacterial MDHs, whose *eLnP* is relatively low compared to the rest of the phylogeny. To improve the probability of predicting the true ancestral sequence, we need to improve the *eLnP* of an ancestral sequence, which can be accomplished by perhaps including more sequence data targeted at this clade or by improving the phylogenetic model.

**Fig. 9:**
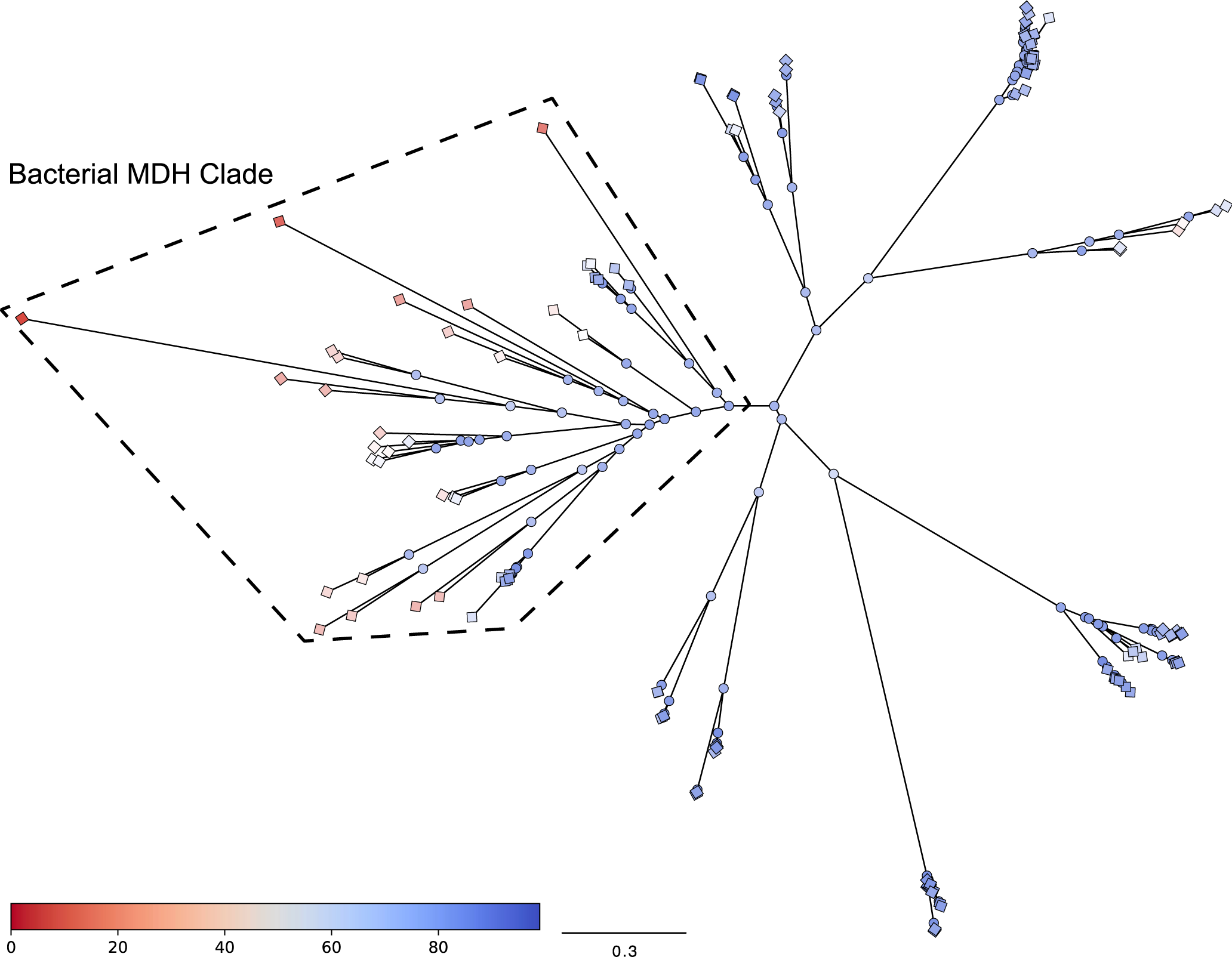
The LG+FO+G12 tree colored by relative total site *eLnP* illustrates the location of uncertain sequences and regions of poor predictiveness. The colored bar in the figure represents the normalized total site *eLnP*; red is the highest *eLnP* and blue is the lowest *eLnP*. Terminal nodes are represented by colored diamonds and internal nodes are represented by colored circles. The bacterial MDH clade is identified by the polygon outlined with dashed lines.

## Conclusion

We have developed a CV method called extant sequence reconstruction, ESR, that can be used to evaluate the accuracy of ASR by comparing its reconstructions against known, true proteins. Using ESR we are able to learn how different phylogenetic models impact reconstructions in three key ways. First, improving a model (as judged by comparing AIC or BIC) should improve the true residue probability. Second, improving the model generally increases the expected chemical similarity between the reconstruction and the true sequence. The two main factors in improving ancestral sequence reconstructions are the choice of substitution matrix and among-site rate variation. Third, the average probability of an SMP reconstruction is a poor judge of the performance of a model. A better model frequently results in lower average probabilities for SMP reconstructions. Taken together, these takeaways point to the importance of performing model selection when using ASR to increase the chances of reconstructing the most accurate sequence.

In this study we compare sequence reconstructions to truth using ESR. However, ESR provides a general framework to “resurrect” extant proteins and experimentally compare their biophysical properties (e.g., melting temperature or activity) to that of the true proteins. An important application of ESR will be in assessing the accuracy of ASR methodology experimentally by comparing the biophysical properties of ESR resurrected proteins to the true modern proteins, a research goal we will address in future work.

## Materials and Methods

### Simulated Datasets

We used INDELible (version 1.03) to generate extant and ancestral datasets. Sequences were simulated under the LG+FO+G12 model of evolution using either the L/MDH, Abl/Src-kinase, or terpene synthase phylogeny. The equilibrium frequencies, alpha parameter for rate-variation, and guide tree were the ML estimates from the L/MDH, Abl/Src-kinase, and terpene synthase phylogenies inferred using the LG+FO+G12 model of evolution. The simulated alignment length was 1000 residues long for each simulation and was repeated 10 times. Ten simulations were repeated with the Poisson+FQ model of evolution and phylogeny for each protein families, resulting in 60 total simulations. ASR and ESR was performed on each simulated dataset using various models of evolution: (1) the Poisson substitution matrix with FQ, (2) the LG substitution matrix with FQ, (3) the LG substitution matrix with FO, and (4) the LG substitution matrix with FO and 12-category gamma G12.

### Biological Datasets

Protein sequences for lactate and malate dehydrogenase (L/MDH), kinase, and terpene synthase homologs were obtained using BLAST searches with the National Center for Biotechnology Information *nr* protein sequence database (Wilson et al. 2015). For each dataset a multiple sequence alignment was generated using MAFFT-LINSI (version 7.487) (Katoh 2002). Sequence and alignment statistics for each protein family are listed in Table 1.

### Phylogenetic inference

Maximum likelihood (ML) phylogenies were inferred using IQ-Tree (version 2.1.1) with various evolutionary models for a given multiple sequence alignment (Bershtein et al. 2008; Nguyen et al. 2015). In order of increasing complexity, the models were: (1) the Poisson substitution matrix with equal equilibrium frequencies (FQ), (2) the LG substitution matrix with FQ, (3) the LG substitution matrix with optimized equilibrium frequencies (FO), (4) the LG substitution matrix with FO and 12-category gamma distributed among-site rate variation (G12), (5) the LG substitution matrix with FO, G12, and a proportion of invariant sites (I), and (6) GTR20 with FO and G12.

### ASR methodology provides a distribution of amino acid states for a reconstructed protein

Here we provide a brief overview of standard ASR methodology for calculating ancestral protein sequences. The ancestral sequence probability distribution for a given internal node, hereafter referred to as the ancestral distribution or simply the “reconstruction”, is an inferred amino acid distribution for every site in an ancestral sequence conditional on a given phylogeny, model of evolution, and corresponding ML model parameters (Songyang et al. 1995; Yang et al. 1995). The ancestral probability distribution reflects our confidence in the presence of each amino acid at every site in the ancestral sequence (Finnigan et al. 2012; Hanson-Smith et al. 2010; Songyang et al. 1995; Yang et al. 1995; Zhang and Nei 1997).

Consider the simple tree on the left-hand side of figure 1, which shows the phylogenetic relationships among three observed, modern sequences (*A*, *B*, and *C*) and an unobserved, ancestral sequence *D*. Since we assume site-independence, when calculating probabilities of the sequence data we can consider one site at a time (Felsenstein 1981; Yang 1994). The joint probability of the amino acids in the four sequences at a site may be written:

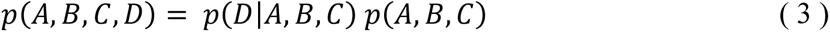

where *p*(*D*|*A*, *B*, *C*) is the probability of observing the amino acid for ancestral sequence *D*, conditional on the observed amino acids in sequences *A*, *B,* and *C*. For clarity we omit the conditional dependencies on the ML phylogeny and model parameters. The probabilities in equation (1) above can be calculated using standard probabilistic models of sequence evolution. In general, we do not know the ancestral states, but the joint probability of the observed amino acids (in sequences *A*, *B*, and *C*) can still be calculated by integrating out the ancestral state in *D* (*i.e.*, by summing over all possible amino acid states at that ancestral site):

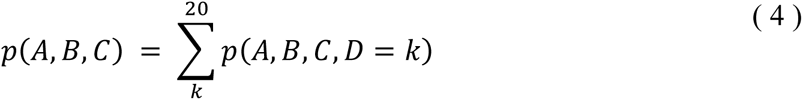

where *k* is one of the possible 20 amino acids (Felsenstein 1981; Yang 1994). This is the usual form of the phylogenetic likelihood function for observed sequence data that is maximized in a ML phylogenetic analysis. In conventional ASR (for the marginal reconstruction), Bayes rule provides the probability distribution for the unobserved ancestral site *D*:

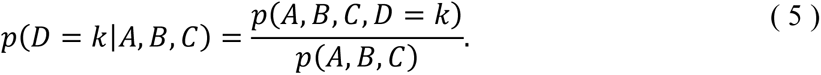

Equation 5 gives the probability that the ancestral state is amino acid *k*. This calculation can be made for every amino acid to produce the full ancestral probability distribution at that site. By repeating the calculation at every site, we construct the full ancestral probability distribution for the ancestral sequence *D* (Songyang et al. 1995; Yang et al. 1995).

The fundamental result of ASR is not a single reconstructed sequence, but rather a distribution that assigns probabilities to all possible sequences of length *N*, the length of the alignment. The extant reconstructed distribution provides all information possible from our evolutionary model and sequence dataset to evaluate the plausibility of different sequences and amino acids at the node of interest. There are two main types of descriptive statistics we can calculate from the reconstructed probability distribution: (1) sequence-specific statistics, which apply only to a single reconstructed sequence (*e.g.*, the SMP), and (2) global reconstruction statistics, which describe an average or expected quantity over all possible reconstructed sequences for a given ancestor. For example, from the reconstructed distribution we can quantify the probability of any specific sequence, sample sequences from the distribution, and generate the SMP sequence (Chang et al. 2002; Eick et al. 2017; Gaucher et al. 2008). Based on the entire distribution, we can calculate the expected average probability and the expected log-probability of the true sequence at the node. In the following we assess the utility of these and other statistics and observe how they are affected by competing evolutionary models of increasing complexity.

### Ancestral and extant reconstructions

Herein we refer to a “reconstruction” as the probability distribution of amino acid states for a single sequence at a hidden node in a phylogeny. For ASR, the hidden node corresponds to an internal node, whereas for extant sequence reconstruction (ESR), the hidden node corresponds to a terminal node. A reconstructed probability distribution is a 20×*N* matrix of *N* sites corresponding to the *N* columns in the sequence alignment. Each site *j* in the reconstruction probability distribution is a categorical distribution represented by a 20-vector of amino acid probabilities that sum to 1. In this work we use what is known as the “marginal reconstruction” of a hidden node (as opposed to the “joint reconstruction”), in which the uncertainties in the hidden states of all other nodes are integrated over (Yang et al. 1995).

### Site-wise CV

The goal of site-wise CV is to phylogenetically reconstruct the sequence of a modern protein. The amino acid probability distribution of a site in a single modern sequence is calculated analogously to how conventional ASR calculates the amino acid probability distribution for an ancestral site. For site-wise CV (Fig. 1, middle right), the training dataset is the original alignment with a single site removed from a single modern sequence corresponding to a terminal node (*i.e.*, a single amino acid was deleted from the alignment). The validation dataset is the single amino acid that is deleted from the original alignment to produce the training dataset.

First, a new ML phylogeny and model parameters were inferred for the training dataset. Depending on the particular phylogenetic model under consideration, the parameters may include one or more of branch lengths, equilibrium frequencies, alpha rate variation, invariant sites fraction, and amino acid exchangeabilities. Then, using the ML parameters from the training set, the probability distribution of the deleted extant amino acid site was calculated. IQ-Tree only reconstructs states for internal nodes of a phylogeny, so a workaround to reconstruct terminal nodes was coded using IQ-Tree and in-house shell and Python scripts. The extant node corresponding to the site and sequence of interest was made an internal node in a new proxy tree by artificially adding two daughter nodes with branch lengths of 1000 to the ML tree constructed from the training set. The corresponding sequence alignment was also modified to include an additional poly-Leu sequence corresponding to one of the new daughter nodes; the other daughter node corresponds to the original sequence of interest with the single site deleted. ASR was then performed with IQ-Tree (by invoking the ‘-asr’ command line option) without optimization using the proxy tree, sequence alignment, and ML parameters previously inferred from the training dataset. This procedure forces IQ-Tree to reconstruct the sequence corresponding to the extant node (now internal) in the proxy tree. Because the two new daughter nodes are attached to the original extant node by very long branches, the daughter nodes contribute no information to the reconstruction at the internal node (a fact which was confirmed empirically by performing analyses with various branch lengths, short to long). These steps were repeated for each site in each sequence in the alignment.

The reconstructed conditional probability distribution for an entire extant sequence was then constructed by collating the conditional probability distribution for each removed site in the extant sequence from the IQ-Tree ASR output files.

### Sequence-wise CV

The purpose of sequence-wise CV is to closely approximate the results of single-site CV by generating a probability distribution for each extant sequence, but with considerably less computation. For sequence-wise CV, the training dataset is the original sequence alignment with no sites removed, and the validation dataset is a single extant sequence. First, using IQ-Tree a maximum likelihood phylogeny was inferred for the alignment that contained the full sequence set. Then, like in the site-wise CV, the original phylogeny was modified to internalize the node corresponding to the chosen extant sequence by making it the parent of two introduced daughter nodes and a poly-Leu sequence was added to the original alignment. Finally, ASR was performed on the modified phylogeny using the ML estimate of each parameter from the original phylogeny to generate a file containing the posterior probability distributions for each internal node. The process was repeated for each sequence in the alignment. A probability distribution is calculated at each site of the alignment, so for sequence-wise CV the corresponding gaps from the true alignment were applied to the extant reconstruction. While this sequence-wise procedure is not strictly CV, the results are extremely close to that of site-wise CV (Supplementary Figure S8) because the respective training sets typically differ by only one residue out of thousands in an entire alignment (e.g., only one difference out of 39,080 total residues for the L/MDH dataset). Sequence-wise CV speeds up the computation time by several orders of magnitude relative to site-wise CV.

To demonstrate that sequence-wise CV reconstruction probabilities are approximately equivalent to site-wise CV probabilities we used the LG+FO+G12 model and the L/MDH dataset we took two approaches. First, we compared the overall true sequence reconstructions between the two CV methods. A line of best fit for the *LnP* of each true sequence reconstruction between the two cross-validation methods has a slope of 1.001 and an intercept of 3.824 (Supplementary Figure S7a). The slope indicates that the true sequence *LnP* from sequence-wise CV approximates closely the true sequence *LnP* from site-wise CV and scales with it almost perfectly. The intercept indicates a small positive offset to the sequence-wise reconstructions, which is expected from the fact that the sequence-wise approximation to *LnP* should always be less than or equal to the correct site-wise value. A histogram of the difference in *LnP* for each true sequence (the difference is sequence-wise *LnP* from site-wise *LnP*) demonstrates that the *LnP* from a site-wise reconstruction is always lower than from a sequence-wise reconstruction (Supplementary Figure S8b). The average difference in *LnP* between two sequences reconstructed from site-wise and sequence-wise is -3.66 with a 0.16 standard error of the mean.

We also compared the individual site reconstructions for the true residues for site-wise CV and sequence-wise CV. A line of best fit for the true residue *LnP* calculated by sequence-wise CV as a function of the true residue *LnP* by site-wise CV is 0.954 and an intercept of -0.007 (Supplementary Figure S8c). Even the individual *LnP* for each sites reconstructed using sequence-wise CV is a good approximation of the site-wise CV. We repeated the process with the probability of each true residue and, like with the *LnP*, sequence-wise CV is a good approximation to site-wise CV (Supplementary Figure S8d). Each site reconstructed requires a new phylogeny and parameters estimated by IQ-Tree, which is 39,080 unique runs for a single alignment. The average absolute error per true residue between the two CV methods is 0.001.

### Phylogenetic tree pruning

To determine if the estimates of fraction correct and of true sequence *LnP* are robust to the branch length, a L/MDH phylogenetic tree was pruned of selected branches and sequence-wise CV was performed. First, for the phylogeny inferred using LG+FO+G12, taxa and adjacent branches were removed, and a list of the remaining taxa was stored. Next, taxa from LG+FO, LG+FQ, and Poisson+FQ were also removed, provided they were removed from the LG+FO+G12 phylogeny. The internal nodes associated with each branch were also removed while maintaining total branch lengths. Finally, sequence-wise CV was performed on each pruned phylogenetic tree. For each dataset the slope was calculated for a line of best fit between average fraction correct against average SMP sequence probability (averaging over all extant reconstructions).

### Single Most Probable (SMP) reconstructed sequence

SMP sequences are generated by selecting the most probable residue at each site from the reconstructed conditional probability distribution for each sequence. Extant SMPs are constructed from an extant reconstruction probability distribution; ancestral SMPs are constructed from an ancestral reconstruction probability distribution. In many publications, SMP sequences are referred to as “maximum likelihood (ML) ancestral sequences”, which strictly is a misnomer because the SMP state is not an ML estimate (Eick et al. 2017; Wheeler et al. 2016). Reconstructed hidden states are not parameters of the likelihood function; rather, they are unobserved data states that are integrated out of the likelihood function.

### Log-probability of a specific sequence

The log-probability (*LnP*) of a sequence is the sum of the *LnP* for the amino acid state *a* at each site *j* of the sequence:

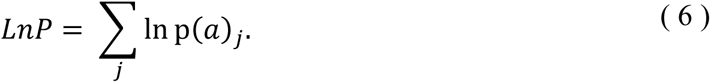

### Expected log-probability for a reconstruction

The expected log-probability (*eLnP*) of a reconstruction can be thought of as the average log-probability of a sequence sampled from the reconstructed distribution:

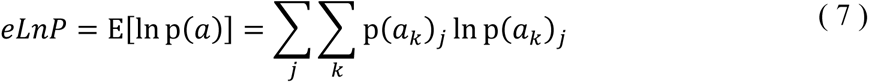

where *ak* is the amino acid state, which can be any one *k* of the 20 amino acids, and *p (ak)j* is the probability of the amino acid state *ak* at site *j* of the sequence. Note that here the expectation is taken over the entire reconstructed probability distribution (*i.e.*, over all sites and over all 20 possible amino acid states). The *eLnP* of a reconstruction is equivalent to the negative entropy of the reconstructed probability distribution and is an estimate of the log-probability of the corresponding true sequence.

### Average probability or expected fraction correct for a specific sequence

For a given sequence *i*, the expected fraction *f* of correct residues is equal to the arithmetic average of the probabilities for the amino acid at each site *j* in the sequence:

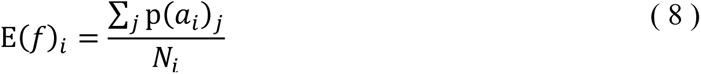

where *Ni* is the length of sequence *i*, and *ai* is the amino acid state at site *j* in sequence *i*. The “fraction correct” *f* is defined as the actual number of correct residues in the sequence divided by the sequence length. We usually refer to the expected fraction correct as simply the “average probability” of a reconstructed sequence.

Because we model the probability of successfully identifying the correct amino acid at each site as an independent Bernoulli, the average probability over sites is a scaled sum of independent Bernoulli probabilities. Hence, both the expected number of correct residues and the expected fraction of correct residues follow from a Poisson-Binomial distribution (Chen and Liu 1997; Wang 1993).

### Expected fraction correct for a reconstruction

The expected fraction of correct amino acids *f* for a reconstruction is calculated as

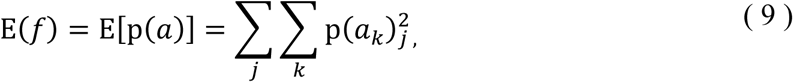

where the summation is taken over sites *j* and amino acids *k*. This expectation can be thought of as the average, over all possible sequences, of the expected fraction correct for each sequence, weighted by the probability of each sequence.

### Chemical similarity between SMP and true sequences

To account for chemical similarity among mutations, we calculated the Grantham distance (*Gd*) between the SMP sequence and the true extant sequence as

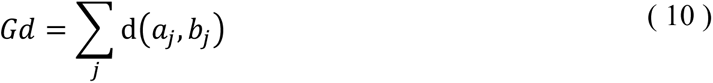

where d (*a*,*b*) is Grantham’s distance between the true amino acid *a* and the SMP amino acid *b*, summing over each site *j* in the sequence (Grantham 1974). To calculate *Gd* for an entire alignment we sum over each protein in the alignment.

### Chemical similarity between a sequence reconstruction and the true sequence

To quantify the expected chemical similarity between the true sequence and sequences sampled from the reconstructed distribution, we calculate an expected *Gd* as

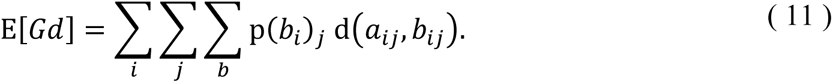

E[*Gd*] is the expected *Gd* and *p (b)* is the probability of the amino acid state. To calculate E[*Gd*] for an entire alignment we sum over all possible amino acids, for each site in a protein, and for each protein in the alignment.

### Model selection information criteria

The Akaike Information Criterion (AIC) and the Bayesian Information Criterion (BIC) were calculated as

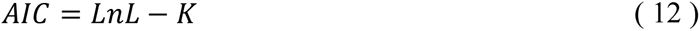

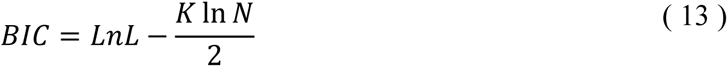

respectively, where *LnL* is the maximum log-likelihood for the model, *K* is the number of free parameters, and *N* is the number of columns in the alignment.

### Tree Visualization

To visualize the uncertainty in our ancestral sequences, we map normalized *eLnP* of each node onto the phylogeny. We calculate the *eLnP* (eqn. 7) for each ancestral and extant node and normalize each ancestral and extant *eLnP* with the maximum and minimum *eLnP*, so that the normalized value is between 0 and 100. Then we modify the Newick formatted tree file so that each node is associated with the corresponding normalized *eLnP*. The subsequent tree was displayed with FigTree (version 1.4.3).

## Acknowledgements

This work was supported by the National Institute for General Medicine at the National Institutes of Health (grant numbers R01 GM096053 and R01 GM132499).

## Code and Data availability

The method developed in this paper utilizes Python and Bash scripting to utilize IQ-Tree to reconstruct extant sequences in addition to ancestral sequences. Extant phylogenies, alignments and code used in ESR is publicly available at https://github.com/sennettm/Extant_Sequence_Reconstruction_ESR.

