## Supplemental for "Extant Sequence Reconstruction: The accuracy of ancestral sequence reconstructions evaluated by extant sequence cross-validation"

### Supplementary Information

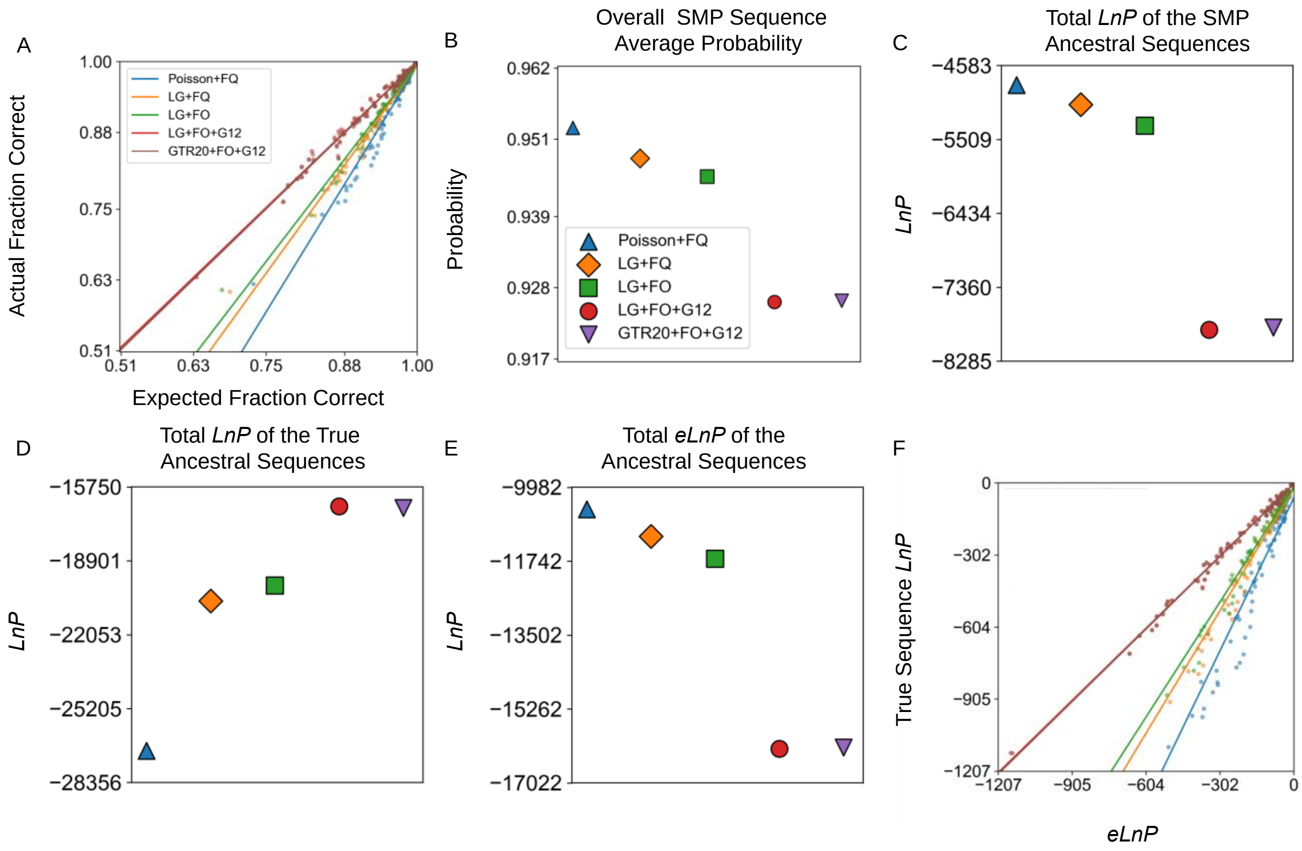

**Fig. S1: ASR probabilities for Abl/Src-kinase simulated sequences are accurate when the model is true or overparameterized.** Ten sets of ancestral sequences were simulated using the LG+FO+G12 model of evolution on a Abl/Src-kinase phylogeny inferred using the same model of evolution. The analyses of ancestral reconstructions for the tenth dataset are shown. (*a*) A plot of the actual fraction correct against expected fraction correct for each reconstructed ancestral sequence in a simulation for each model of evolution. The corresponding line of best fit is shown. (*b*) Each SMP sequence has an average probability. We plot the average of all average SMP sequence probabilities for each model of evolution. (*c*-*e*) The total *LnP* of all SMP sequences, true sequences, and *eLnP* for each model of evolution. (*f*) A plot of true sequence *LnP* vs *eLnP* for each reconstructed ancestral sequence for each model of evolution. The slopes for (*a*) and (*f*) are given in Supplementary Table 2. The values for (*c*-*e*) are given in Supplementary Table 8.

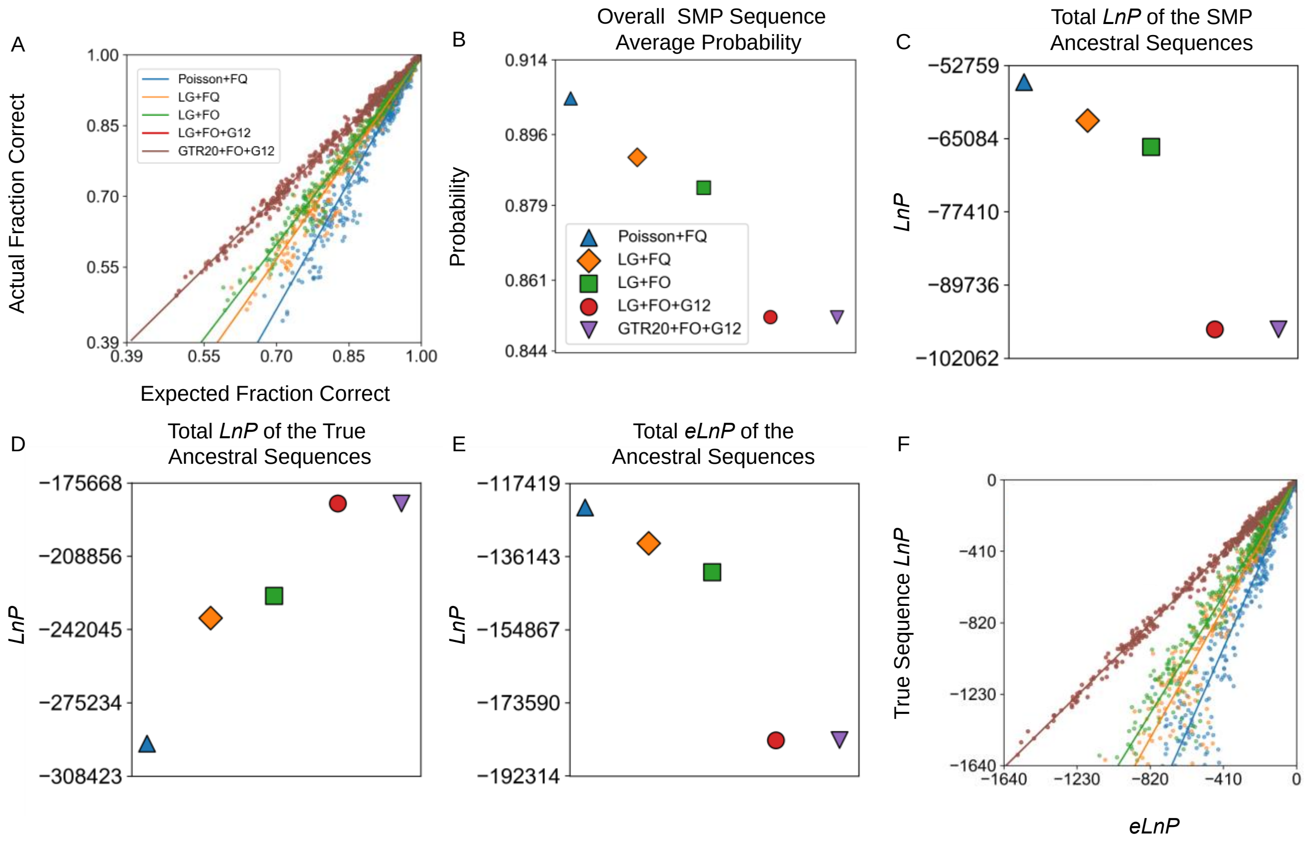

**Fig. S2: ASR probabilities for terpene synthase simulated sequences are accurate when the model is true or overparameterized.** Sequences were simulated under the LG+FO+G12 model of evolution using the terpene synthase phylogeny and parameter estimates. *a*-*f* are the same as in Supplementary Figure 1*a*-*f*. The slopes for (*a*) and (*f*) are given in Supplementary Table 2. The values for (*c*-*e*) are given in Supplementary Table 8.

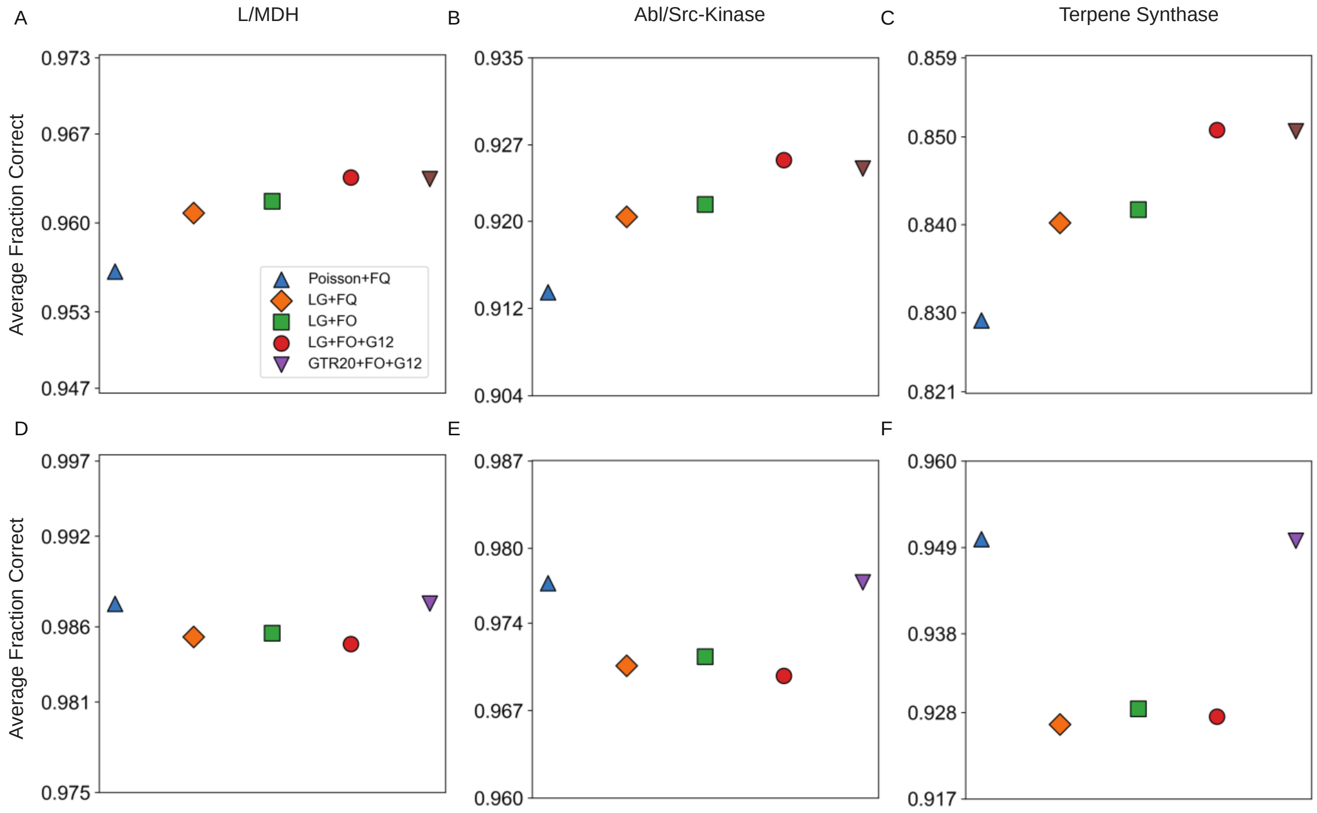

**Fig. S3: Average fraction correct of ASR improves as the model is more correctly specified.** (*a*-*c*) Average fraction correct for the SMP ancestral sequence for L/MDHs, Abl/Src-Kinases, and terpene synthase protein families. Sequences were simulated using the LG+FO+G12 model of evolution. (*d*-*f*) are the same as (*a*-*c*), but sequences were simulated using the Poisson+FQ model of evolution.

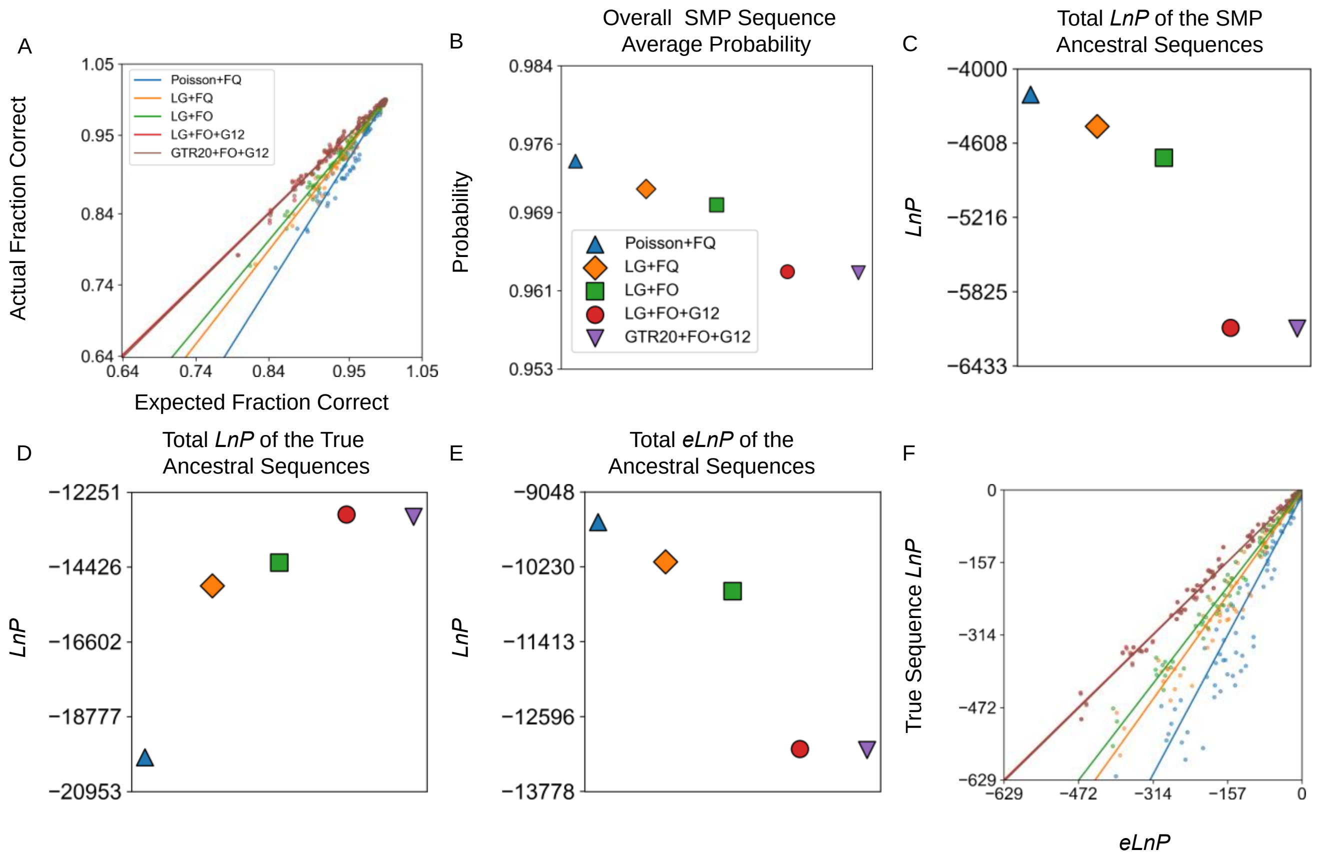

**Fig. S4: ASR probabilities for LMDH simulated sequences are accurate when the model is true or overparameterized.** Sequences were simulated under the LG+FO+G12 model of evolution using the the ML LMDH tree topology, ML branch lengths, and ML model parameters previously estimated with the LG+FO+G12 model. Then ASR was performed with five different evolutionary models using the “true” tree topology and branch lengths while all other parameters were estimated. *a-f* are the same as in Supplementary Figure 1*a-f*. The slopes for (*a*) and (*f*) are given in Supplementary Table 4. The values for (*c-e*) are given in Supplementary Table 9. Note that this figure and analysis is nearly identical to the one shown in Fig. 1 of the main text, except that here the branch lengths were not inferred during phylogenetic analysis and ASR; rather, the branch lengths were fixed to their true values (*i.e.*, the branch lengths used during simulation of the sequences).

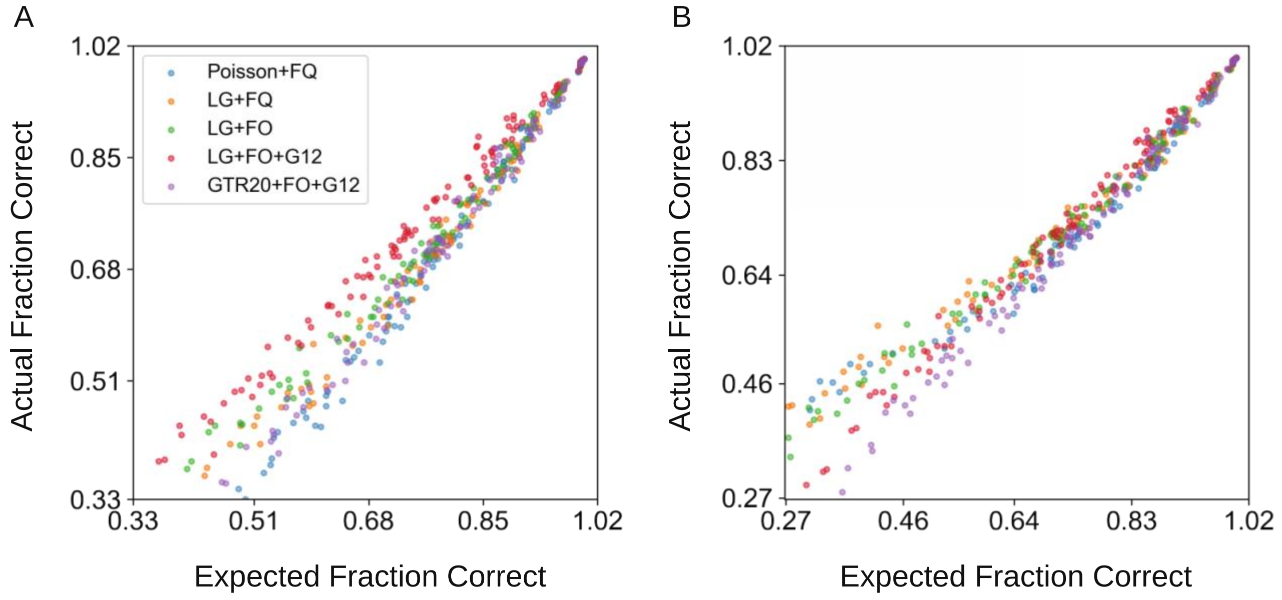

**Fig. S5: For a small three-taxa tree, ASR probabilities for simulated sequences are accurate when the model is true or when using the true branch lengths.** Sequences were simulated along a known tree under the LG+FO+G12 model of evolution using the ML model parameter estimates from the real LMDH dataset in Fig. 4a. (*a*) A plot of the actual fraction correct against expected fraction correct for each reconstructed ancestral sequence for five models of evolution while allowing all branch lengths to be estimated. (*b*) The same as (*a*), except all branch lengths are set to their true values (*i.e.*, the values of the tree used when simulating the sequences). The slopes for (*a*) and (*b*) are given in Supplementary Table 5. Note that these two analyses are similar to that shown in main text Fig 1a and Supplemtary Fig S4a, except that the tree used in these simulations has only three taxa.

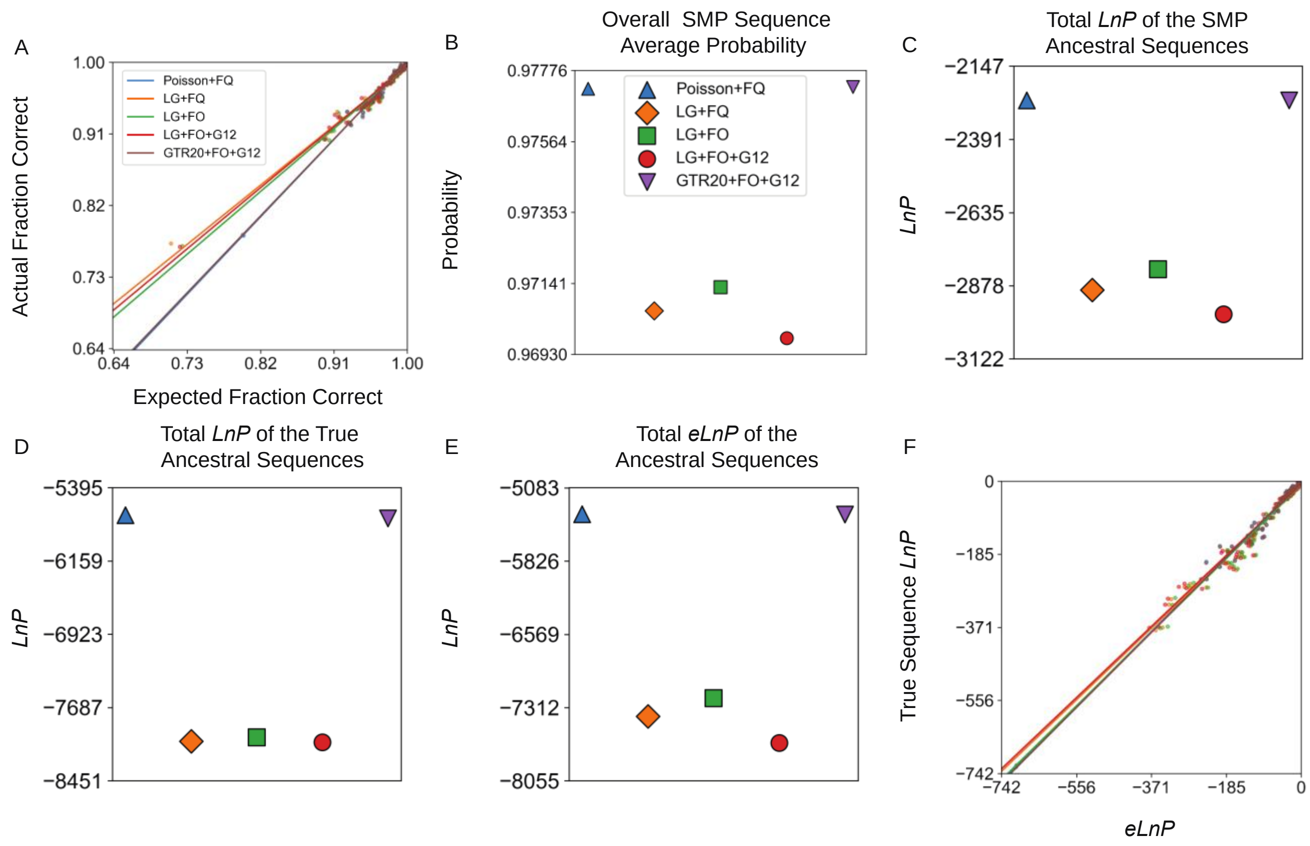

**Fig. S6: ASR probabilities for Abl/Src-kinase simulated sequences are accurate even when the model is misspecified.** Ten sets of ancestral sequences were simulated using the Poisson+FQ model of evolution on a Abl/Src-kinase phylogeny inferred using the same model of evolution. *a*-*f* are the same as in Supplementary Figure 1*a*-*f*. The slopes for (*a*) and (*f*) are given in Supplementary Table 3. The values for (*c*-*e*) are given in Supplementary Table 10.

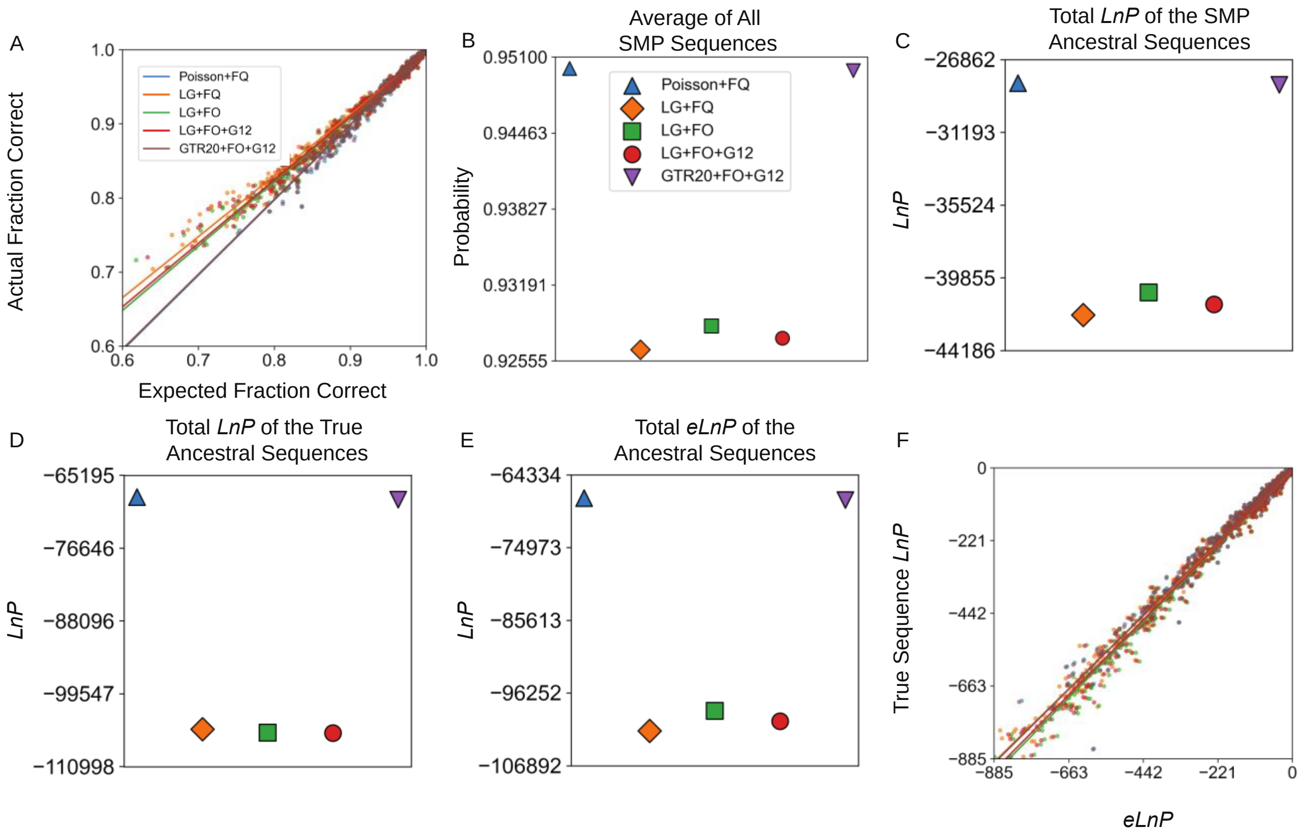

**Fig. S7: ASR probabilities for terpene synthase simulated sequences are accurate even when the model is misspecified.** Ten sets of ancestral sequences were simulated using the Poisson+FQ model of evolution on a terpene synthase phylogeny inferred using the same model of evolution. *a*-*f* are the same as in Supplementary Figure 1*a*-*f*. The slopes for (*a*) and (*f*) are given in Supplementary Table 3. The values for (*c*-*e*) are given in Supplementary Table 10.

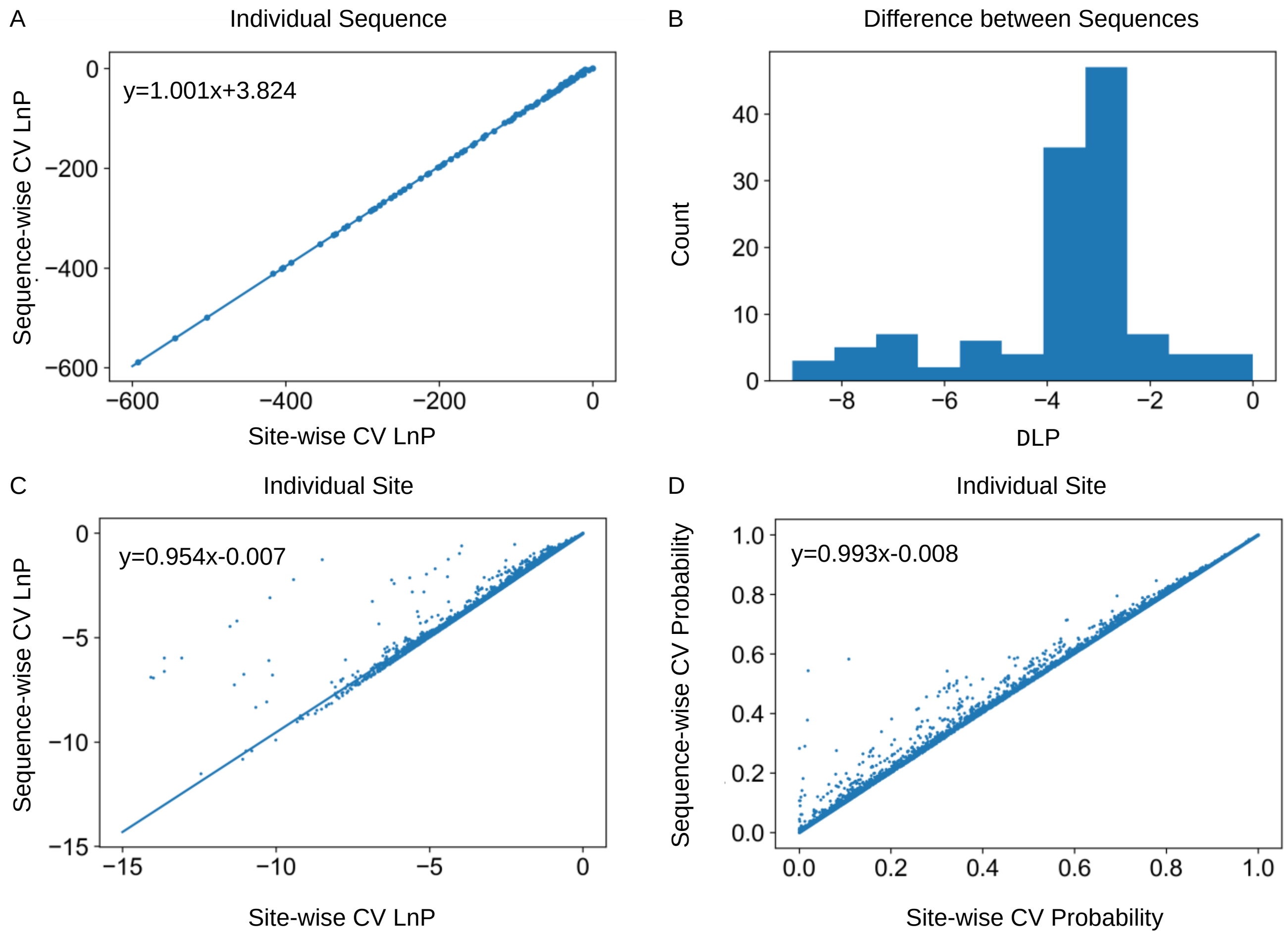

**Fig. S8: True residue probabilities calculated by sequence-wise CV approximate those calculated by site-wise CV.** (*a*) The *LnP* of the true sequence from site-wise and sequence-wise CV. (*b*) The difference in the *LnP* between site-wise and sequence-wise CV for each true sequence. (*c*) The *LnP* of the true residue for sequence-wise CV plotted as a function of site-wise CV. (*d*) Same as in (*c*), except with the probability rather than log-probability.

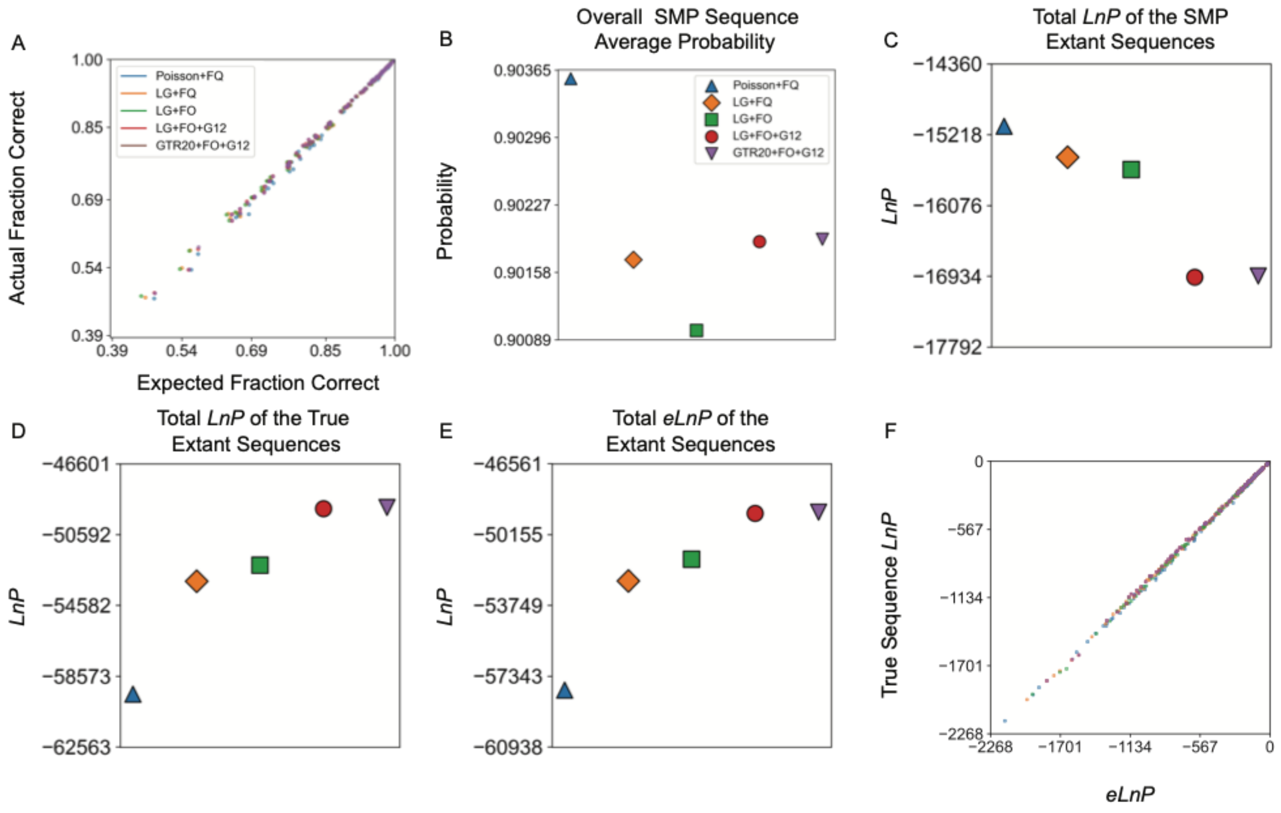

**Fig. S9: ESR probabilities for extant L/MDH sequences simulated under a LG+FO+G12 model are accurate when the model is misspecified.** Analyses on the extant reconstructions for the tenth dataset of L/MDH sequences simulated under the LG+FO+G12 model of evolution are shown above. *a*-*f* are the same as in Supplementary Figure 2*a*-*f*. The slopes for (*a*) and (*f*) are given in Supplementary Table 6. The values for (*c*-*e*) are given in Supplementary Table 11.

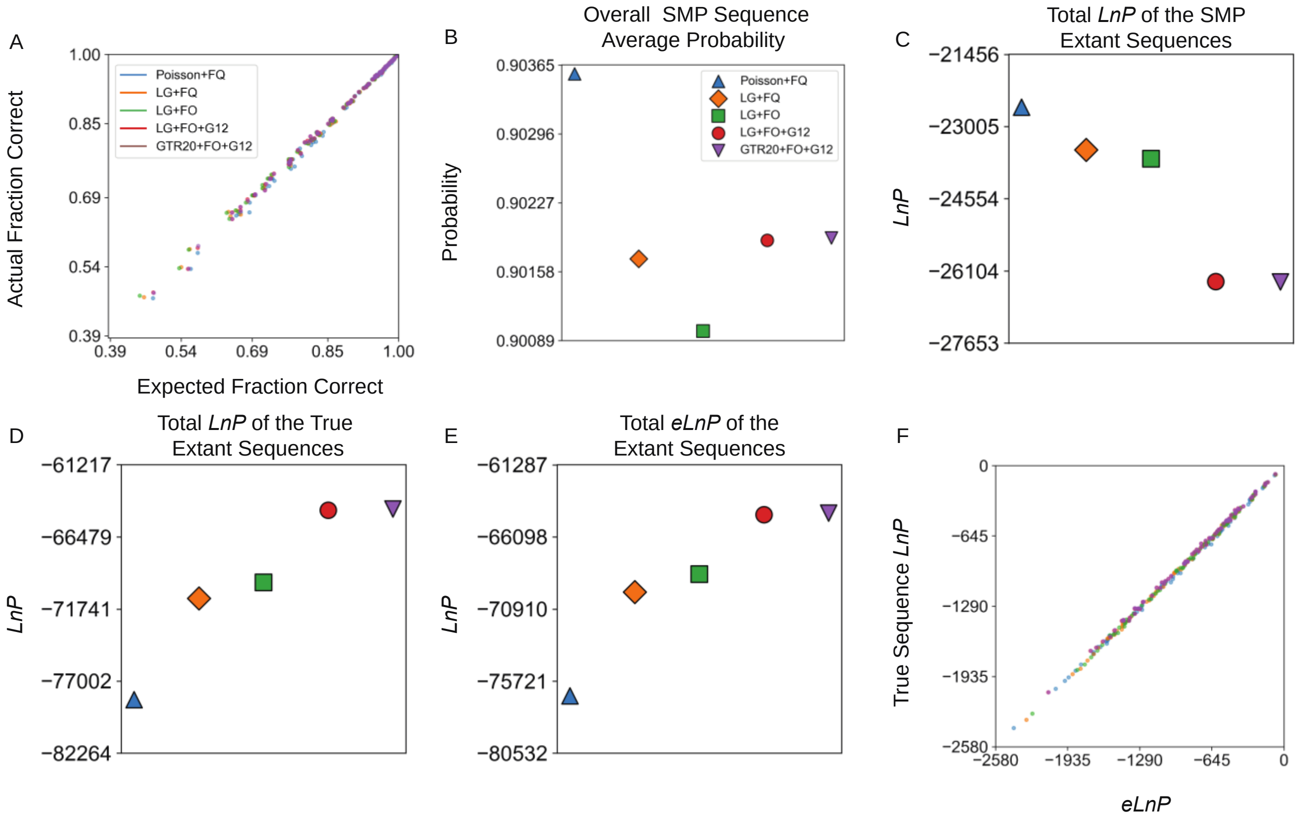

**Fig. S10: ESR probabilities for extant Abl/Src-kinase sequences simulated under a LG+FO+G12 model are accurate when the model is misspecified.** Analyses on the extant reconstructions for the tenth dataset of Abl/Src-kinase sequences simulated under the LG+FO+G12 model of evolution are shown above. *a*-*f* are the same as in Supplementary Figure 2*a*-*f*. The slopes for (*a*) and (*f*) are given in Supplementary Table 6. The values for (*c*-*e*) are given in Supplementary Table 11.

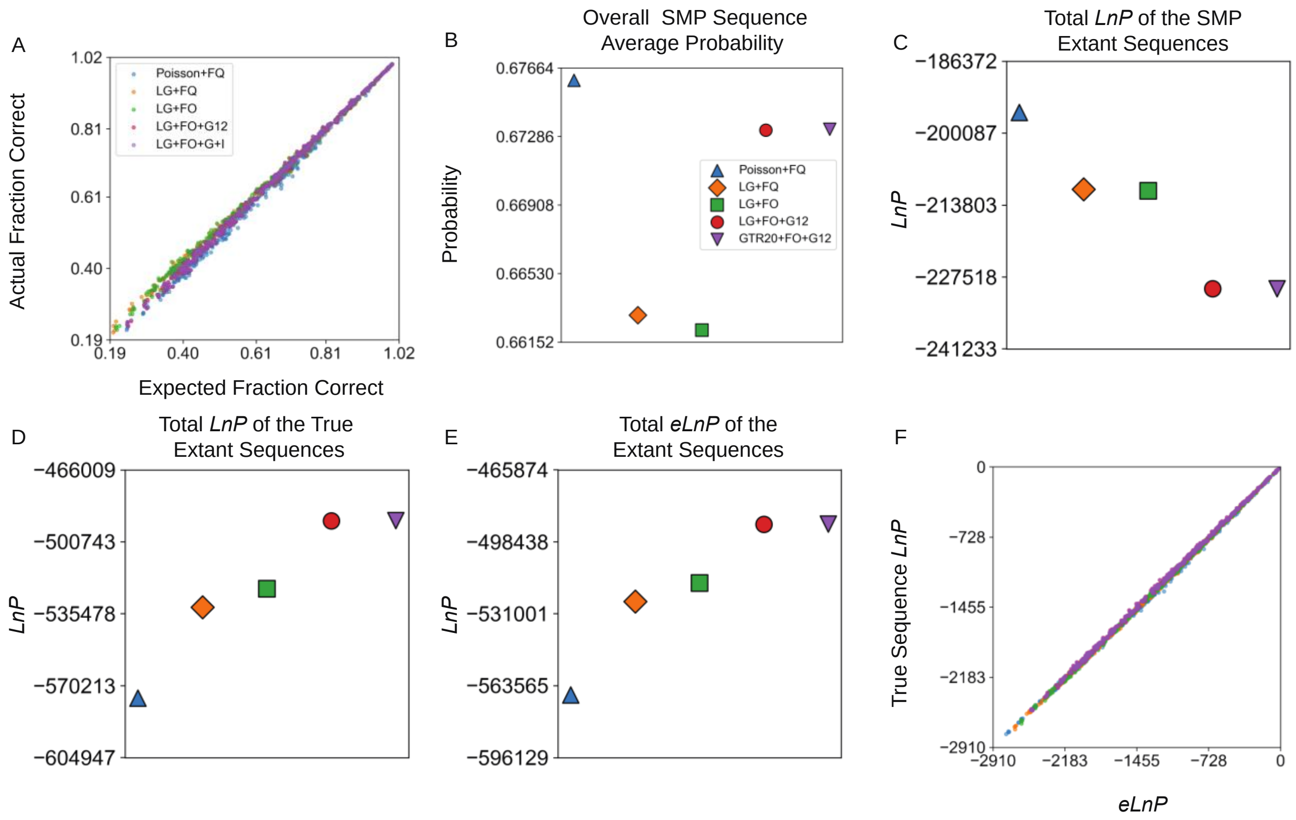

**Fig. S11: ESR probabilities for extant terpene synthase sequences simulated under a LG+FO+G12 model are accurate when the model is misspecified.** Analyses on the extant reconstructions for the tenth dataset of terpene synthase sequences simulated under the LG+FO+G12 model of evolution are shown above. *a*-*f* are the same as in Supplementary Figure 2*a*-*f*. The slopes for (*a*) and (*f*) are given in Supplementary Table 6. The values for (*c*-*e*) are given in Supplementary Table 11.

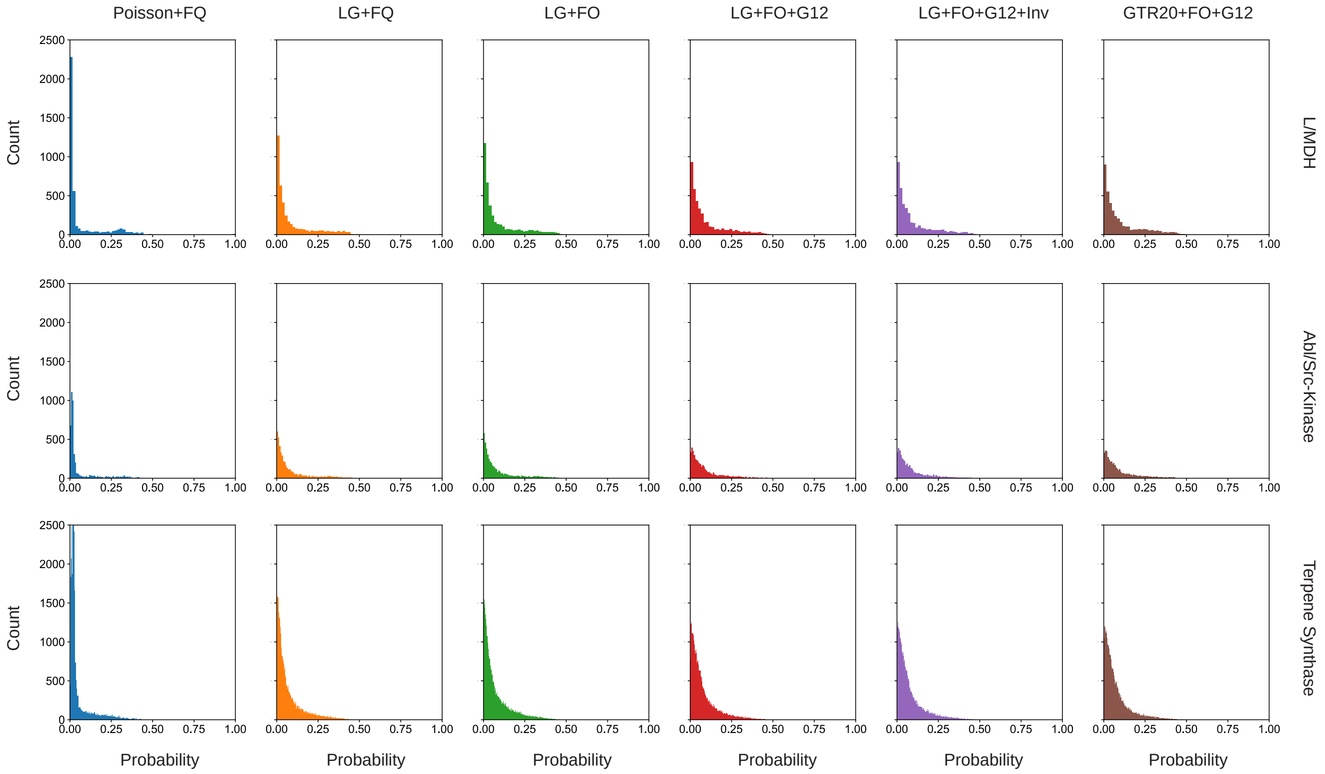

**Fig. S12: Increasing model complexity increases the true residue probability.** Distributions of true residue posterior probabilities at incorrect sites for different models of evolution and protein families.

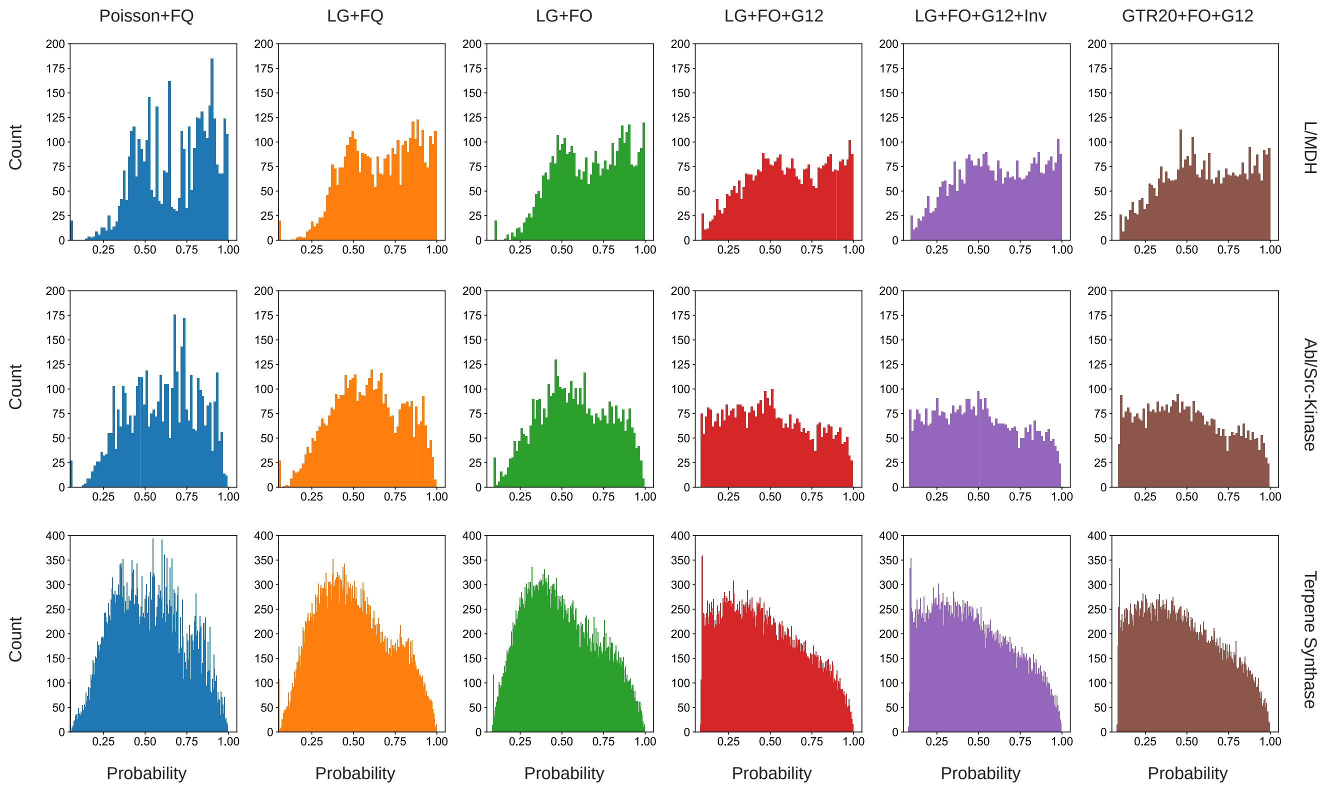

**Fig. S13: Increasing model complexity decreases the probability of the incorrect SMP residue.** Distributions of SMP residue posterior probabilities at incorrect sites for different models of evolution and protein families.

| Real L/MDH (ESR) | | | | |
|---|---|---|---|---|
| Model | true *LnP* vs. *eLnP* | | fraction correct vs. <PP> | |
| | slope | intercept | slope | intercept |
| Poisson+FQ | 1.030 | -3.446 | 1.057 | -0.057 |
| LG+FQ | 1.024 | -2.405 | 1.025 | -0.027 |
| LG+FO | 1.025 | -2.184 | 1.010 | -0.013 |
| LG+FO+G12 | 1.004 | -1.588 | 1.039 | -0.039 |
| LG+FO+G12+I | 1.000 | -1.717 | 1.029 | -0.030 |
| GTR20+FO+G12 | 1.006 | 1.931 | 1.023 | -0.025 |
| Real Abl/Src-Kinase (ESR) | | | | |
| Model | true *LnP* vs. *eLnP* | | fraction correct vs. <PP> | |
| | slope | intercept | slope | intercept |
| Poisson+FQ | 0.997 | -11.636 | 1.042 | -0.050 |
| LG+FQ | 0.997 | -7.161 | 0.987 | 0.002 |
| LG+FO | 0.992 | -7.608 | 0.980 | 0.009 |
| LG+FO+G12 | 1.004 | -1.663 | 1.025 | -0.029 |
| LG+FO+G12+I | 0.998 | -1.973 | 1.016 | -0.021 |
| GTR20+FO+G12 | 1.015 | 0.186 | 1.017 | -0.020 |
| Real Terpene Synthase (ESR) | | | | |
| Model | true *LnP* vs. *eLnP* | | fraction correct vs. <PP> | |

| | slope | intercept | slope | intercept |
|---|---|---|---|---|
| Poisson+FQ | 1.009 | -8.815 | 1.045 | -0.051 |
| LG+FQ | 1.012 | -8.721 | 0.994 | 0.004 |
| LG+FO | 1.007 | -8.419 | 0.995 | 0.008 |
| LG+FO+G12 | 1.012 | -2.147 | 1.010 | -0.011 |
| LG+FO+G12+I | 1.011 | -2.282 | 1.008 | -0.010 |
| GTR20+FO+G12 | 1.010 | -0.282 | 1.018 | -0.019 |

SI Table 1: Linear regression slope and intercept summary of ESR for (1) each true extant *LnP* vs the extant *eLnP* and for (2) fraction correct vs average probability of the SMP sequence for each model and dataset from figure 4*a*-*c* and 7*d*.

| Simulated L/MDH (ASR) | | | | |
|---|---|---|---|---|
| Model | true *LnP* vs. *eLnP* | | fraction correct vs. <PP> | |
| | slope | intercept | slope | intercept |
| Poisson+FQ | 2.45 (0.09) | -7.04 (3.18) | 1.78 (0.05) | -0.78 (0.05) |
| LG+FQ | 1.73 (0.05) | -1.86 (2.11) | 1.51 (0.04) | -0.51 (0.05) |
| LG+FO | 1.54 (0.05) | -0.87 (1.71) | 1.38 (0.03) | -0.38 (0.03) |
| LG+FO+G12 | 1.01 (0.02) | -0.25 (0.98) | 1.01 (0.01) | -0.01 (0.01) |
| GTR20+FO+G12 | 1.02 (0.02) | -0.37 (1.01) | 1.02 (0.01) | -0.02 (0.01) |
| Simulated Abl/Src-Kinase (ASR) | | | | |
| Model | true *LnP* vs. *eLnP* | | fraction correct vs. <PP> | |
| | slope | intercept | slope | intercept |
| Poisson+FQ | 2.26 (0.12) | -46.45 (12.44) | 1.72 (0.07) | -0.72 (0.07) |
| LG+FQ | 1.74 (0.07) | -14.40 (5.66) | 1.45 (0.04) | -0.45 (0.04) |
| LG+FO | 1.63 (0.06) | -10.07 (5.12) | 1.37 (0.04) | -0.38 (0.04) |
| LG+FO+G12 | 1.01 (0.02) | 0.37 (3.89) | 1.01 (0.02) | -0.01 (0.02) |
| GTR20+FO+G12 | 1.02 (0.02) | -0.28 (4.07) | 1.01 (0.02) | -0.01 (0.02) |

| | | | | |
|---|---|---|---|---|
| Simulated Terpene Synthase (ASR) | | | | |
| Model | true *LnP* vs. *eLnP* | | fraction correct vs. <PP> | |
| | slope | intercept | slope | intercept |
| Poisson+FQ | 2.29 (0.02) | -23.71 (4.67) | 1.78 (0.01) | -0.78 (0.01) |
| LG+FQ | 1.89 (0.02) | 26.34 (5.27) | 1.47 (0.02) | -0.47 (0.01) |
| LG+FO | 1.70 (0.02) | 23.11 (4.95) | 1.35 (0.01) | -0.35 (0.01) |
| LG+FO+G12 | 1.00 (0.01) | -2.49 (2.53) | 1.01 (0.01) | -0.01 (0.01) |
| GTR20+FO+G12 | 1.00 (0.01) | -2.62 (2.61) | 1.01 (0.01) | -0.01 (0.01) |

SI Table 2: Linear regression slope and intercept summary of ASR for (1) each true ancestral *LnP* vs the ancestral *eLnP* and for (2) fraction correct vs average probability of the SMP sequence for each model and dataset from figure 1*a*,*f* and Supplementary Figure 1*a*,*f* and 2*a*,*f* with data simulated using the LG+FO+G12 evolutionary model.

| Simulated L/MDH (ASR) | | | | |
|---|---|---|---|---|
| Model | true *LnP* vs. *eLnP* | | fraction correct vs. <PP> | |
| | slope | intercept | slope | intercept |
| Poisson+FQ | 1.01 (0.03) | -0.19 (0.70) | 1.01 (0.03) | -0.01 (0.03) |
| LG+FQ | 1.04 (0.03) | -2.23 (0.74) | 0.96 (0.03) | 0.04 (0.03) |
| LG+FO | 1.07 (0.03) | -2.05 (0.73) | 0.99 (0.03) | 0.01 (0.03) |
| LG+FO+G12 | 1.02 (0.03) | -1.95 (0.73) | 0.95 (0.03) | 0.05 (0.03) |
| GTR20+FO+G12 | 1.01 (0.03) | -0.26 (0.70) | 1.02 (0.03) | -0.02 (0.03) |
| Simulated Abl/Src-Kinase (ASR) | | | | |
| Model | true *LnP* vs. *eLnP* | | fraction correct vs. <PP> | |
| | slope | intercept | slope | intercept |
| Poisson+FQ | 1.00 (0.02) | -0.83 (1.90) | 1.01 (0.04) | -0.02 (0.04) |
| LG+FQ | 0.97 (0.03) | -8.95 (3.26) | 0.83 (0.03) | 0.16 (0.03) |
| LG+FO | 1.00 (0.03) | -8.24 (3.33) | 0.87 (0.03) | 0.13 (0.03) |
| LG+FO+G12 | 0.96 (0.03) | -6.65 (3.23) | 0.84 (0.03) | 0.16 (0.03) |
| GTR20+FO+G12 | 1.01 (0.03) | -0.97 (2.00) | 1.02 (0.04) | -0.02 (0.04) |

| | | | | |
|---|---|---|---|---|
| Simulated Terpene Synthase (ASR) | | | | |
| Model | true *LnP* vs. *eLnP* | | fraction correct vs. <PP> | |
| | slope | intercept | slope | intercept |
| Poisson+FQ | 1.00 (0.01) | -0.89 (1.54) | 1.01 (0.01) | -0.07 (0.01) |
| LG+FQ | 0.99 (0.01) | -8.09 (1.72) | 0.83 (0.05) | 0.16 (0.05) |
| LG+FO | 1.04 (0.01) | -5.65 (1.58) | 0.88 (0.01) | 0.12 (0.01) |
| LG+FO+G12 | 1.02 (0.01) | -5.47 (1.59) | 0.86 (0.01) | 0.13 (0.01) |
| GTR20+FO+G12 | 1.00 (0.01) | -1.01 (1.54) | 1.01 (0.01) | -0.01 (0.01) |

SI Table 3: Linear regression slope and intercept summary of ASR (as in SI Table 2). Here slopes and intercepts are from figure 2*a*,*f* and Supplementary Figure 6*a*,*f* and 7*a*,*f* with data simulated using the Poisson+FQ evolutionary model.

| Simulated LMDH (ASR with true tree) | | | | |
|---|---|---|---|---|
| Model | true *LnP* vs. *eLnP* | | fraction correct vs. <PP> | |
| | slope | intercept | slope | intercept |
| Poisson+FQ | 1.94 (0.05) | -16.09 (2.61) | 1.61 (0.04) | -0.62 (0.04) |
| LG+FQ | 1.40 (0.03) | -6.23 (1.72) | 1.30 (0.02) | -0.30 (0.02) |
| LG+FO | 1.32 (0.02) | -3.82 (1.52) | 1.22 (0.01) | -0.22 (0.01) |
| LG+FO+G12 | 1.00 (0.01) | 0.38 (1.02) | 1.00 (0.01) | 0.00 (0.01) |
| GTR20+FO+G12 | 1.01 (0.01) | 0.30 (1.08) | 1.00 (0.01) | 0.00 (0.01) |

SI Table 4: Linear regression slope and intercept summary of ASR for (1) each true ancestral *LnP* vs the ancestral *eLnP* and for (2) fraction correct vs average probability of the SMP sequence for each model and dataset from Supplementary Figure 4*a*,*f* with data simulated using the LG+FO+G12 evolutionary model and reconstructed using the true tree topology and branch lengths

| Simulated LMDH (ASR) | | | | |
|---|---|---|---|---|
| Model | fraction correct vs. <PP> ML branch | | fraction correct vs. <PP> True branch | |
| | slope | intercept | slope | intercept |
| Poisson+FQ | 1.31 | -0.32 | 0.72 | 0.19 |
| LG+FQ | 1.14 | -0.16 | 0.81 | 0.15 |
| LG+FO | 1.06 | -0.09 | 0.87 | 0.10 |
| LG+FO+G12 | 0.99 | 0.00 | 0.99 | 0.00 |
| GTR20+FO+G12 | 1.23 | -0.24 | 1.07 | -0.08 |

SI Table 5: Linear regression slope and intercept summary of ASR for fraction correct vs average probability of the SMP sequence for each model from Supplementary Figure 5*a,b* with data simulated using the LG+FO+G12 evolutionary model and reconstructed using ML branch lengths or true branch lengths

| Simulated L/MDH (ESR) | | | | |
|---|---|---|---|---|
| Model | true *LnP* vs. *eLnP* | | fraction correct vs. <PP> | |
| | slope | intercept | slope | intercept |
| Poisson+FQ | 1.018 | -3.867 | 1.029 | -0.029 |
| LG+FQ | 1.013 | -0.306 | 0.991 | -0.008 |
| LG+FO | 1.018 | 0.096 | 0.981 | -0.018 |
| LG+FO+G12 | 1.006 | 2.053 | 1.001 | -0.001 |
| GTR20+FO+G12 | 1.006 | 2.004 | 1.003 | -0.003 |
| Simulated Abl/Src-Kinase (ESR) | | | | |
| Model | true *LnP* vs. *eLnP* | | fraction correct vs. <PP> | |
| | slope | intercept | slope | intercept |
| Poisson+FQ | 0.994 | -27.456 | 1.016 | -0.019 |
| LG+FQ | 1.010 | -5.995 | 0.959 | 0.034 |
| LG+FO | 1.011 | -5.682 | 0.950 | 0.042 |
| LG+FO+G12 | 0.997 | -1.856 | 1.0008 | -0.0002 |
| GTR20+FO+G12 | 0.997 | -1.447 | 0.9992 | 0.0003 |
| Simulated Terpene Synthase (ESR) | | | | |
| Model | true *LnP* vs. *eLnP* | | fraction correct vs. <PP> | |
| | slope | intercept | slope | intercept |

| | | | | |
|---|---|---|---|---|
| Poisson+FQ | 0.998 | -22.970 | 1.023 | -0.025 |
| LG+FQ | 1.006 | -8.354 | 0.961 | 0.032 |
| LG+FO | 1.007 | -7.096 | 0.961 | 0.033 |
| LG+FO+G12 | 1.002 | 2.575 | 1.003 | -0.002 |
| GTR20+FO+G12 | 1.002 | 2.528 | 1.002 | -0.002 |

SI Table 6: Linear regression slope and intercept summary of ESR (as in SI Table 1). Here slopes and intercepts are from Supplementary Figures 9*a*,*f*, 10*a*,*f*, and 11*a*,*f* with data simulated using the LG+FO+G12 evolutionary model.

| Real L/MDH (ESR) | | | |
|---|---|---|---|
| Model | true *LnP* | *eLnP* | SMP *LnP* |
| Poisson+FQ | -18662 | -17696 | -4644 |
| LG+FQ | -16513 | -15840 | -4749 |
| LG+FO | -16279 | -15624 | -4781 |
| LG+FO+G12 | -15388 | -15139 | -5139 |
| LG+FO+G12+I | -15377 | -15171 | -5164 |
| GTR20+FO+G12 | -15172 | -14859 | -5203 |
| Real Abl/Src-Kinase (ESR) | | | |
| Model | true *LnP* | e*LnP* | SMP *LnP* |
| Poisson+FQ | -19707 | -18889 | -5514 |
| LG+FQ | -17799 | -17305 | -5870 |
| LG+FO | -17467 | -17025 | -5951 |
| LG+FO+G12 | -16283 | -16066 | -6586 |
| LG+FO+G12+I | -16278 | -16145 | -6593 |
| GTR20+FO+G12 | -16105 | -15864 | -6628 |
| Real Terpene Synthase (ESR) | | | |
| Model | true *LnP* | *eLnP* | SMP *LnP* |
| Poisson+FQ | -160472 | -155396 | -51256 |
| LG+FQ | -148919 | -143606 | -54918 |
| LG+FO | -146168 | -141627 | -55233 |
| LG+FO+G12 | -141998 | -139489 | -58822 |
| LG+FO+G12+I | -141983 | -139503 | -58842 |
| GTR20+FO+G12 | -141236 | -139714 | -57911 |

SI Table 7: Summary of the total *LnP* for the true sequence from figure 5*b*, total *LnP* for the SMP sequences from figure 5*a*, and the total expected *LnP* for each model of evolution and protein families from figure 7*a*.

| Simulated L/MDH (ASR) | | | |
|---|---|---|---|
| Model | true *LnP* | *eLnP* | SMP *LnP* |
| Poisson+FQ | -21565 (810) | -8453 (394) | -3922 (181) |
| LG+FQ | -15738 (516) | -8974 (436) | -3960 (213) |
| LG+FO | -14873 (475) | -9568 (463) | -4244 (222) |
| LG+FO+G12 | -12968 (451) | -12807 (583) | -5932 (301) |
| GTR20+FO+G12 | -13028 (452) | -12788 (595) | -5922 (310) |
| Simulated Abl/Src-Kinase (ASR) | | | |
| Model | true *LnP* | e*LnP* | SMP *LnP* |
| Poisson+FQ | -25782 (1489) | -9892 (586) | -4541 (248) |
| LG+FQ | -19340 (1097) | -10534 (639) | -4754 (306) |
| LG+FO | -18680 (1064) | -11034 (666) | -4998 (319) |
| LG+FO+G12 | -15601 (918) | -15454 (870) | -7458 (484) |
| GTR20+FO+G12 | -15670 (929) | -15405 (883) | -7429 (496) |
| Simulated Terpene Synthase (ASR) | | | |
| Model | true *LnP* | *eLnP* | SMP *LnP* |
| Poisson+FQ | -308215 (10522) | -130364 (4993) | -58602 (2267) |
| LG+FQ | -251346 (9520) | -139075 (5311) | -65087 (2636) |
| LG+FO | -240820 (9004) | -147056 (5709) | -69981 (2929) |
| LG+FO+G12 | -194965 (7015) | -193193 (7140) | -103172 (4184) |
| GTR20+FO+G12 | -195070 (7003) | -193167 (7148) | -103174 (4155) |

SI Table 8: Summary of total *LnP* as in Supplementary Table 7, but with ancestral data simulated using the LG+FO+G12 evolutionary model from figure 1*c*-*e* and Supplementary Figures 1*c*-*e* and 2*c*-*e*.

| Simulated LMDH (ASR using true tree) | | | |
|---|---|---|---|
| Model | true *LnP* | *eLnP* | SMP *LnP* |
| Poisson+FQ | -20133 (813) | -9356 (366) | -4099 (176) |
| LG+FQ | -14881 (539) | -10057 (360) | -4391 (192) |
| LG+FO | -14275 (489) | -10471 (369) | -4633 (184) |
| LG+FO+G12 | -12870 (449) | -12925 (528) | -5994 (269) |
| GTR20+FO+G12 | -12929 (449) | -12903 (537) | -5981 (276) |

SI Table 9: Summary of total *LnP* as in Supplementary Table 7, but with ancestral data simulated using the LG+FO+G12 evolutionary model from Supplementary Figures 4*c*-*e*.

| Simulated L/MDH (ASR) | | | |
|---|---|---|---|
| Model | true *LnP* | *eLnP* | SMP *LnP* |
| Poisson+FQ | -4691 (78) | -4639 (95) | -2027 (45) |
| LG+FQ | -6658 (130) | -6120 (144) | -2330 (56) |
| LG+FO | -6644 (126) | -5984 (136) | -2284 (56) |
| LG+FO+G12 | -6690 (121) | -6354 (145) | -2406 (61) |
| GTR20+FO+G12 | -4736 (82) | -4660 (106) | -2023 (46) |
| Simulated Abl/Src-Kinase (ASR) | | | |
| Model | true *LnP* | e*LnP* | SMP *LnP* |
| Poisson+FQ | -5517 (105) | -5448 (101) | -2301 (60) |
| LG+FQ | -7955 (222) | -7514 (125) | -2936 (50) |
| LG+FO | -7933 (222) | -7336 (129) | -2867 (53) |
| LG+FO+G12 | -7982 (219) | -7789 (155) | -3025 (66) |
| GTR20+FO+G12 | -5572 (96) | -5469 (103) | -2300 (60) |
| Simulated Terpene Synthase (ASR) | | | |
| Model | true *LnP* | *eLnP* | SMP *LnP* |
| Poisson+FQ | -68379 (1093) | -67822 (897) | -28339 (394) |
| LG+FQ | -103517 (1486) | -101189 (1575) | -41763 (754) |
| LG+FO | -104009 (1470) | -98074 (1569) | -40347 (737) |
| LG+FO+G12 | -104013 (1468) | -99626 (1570) | -4106 (739) |
| GTR20+FO+G12 | -68671 (1102) | -68057 (929) | -28386 (403) |

SI Table 10: Summary of total *LnP* as in Supplementary Table 7, but with ancestral data simulated using the Poisson+FQ evolutionary model from figure 2*c*-*e* and Supplementary Figures 6*c*-*e* and 7*c*-*e*.

| Simulated L/MDH (ESR) | | | |
|---|---|---|---|
| Model | true *LnP* | *eLnP* | SMP *LnP* |
| Poisson+FQ | -59584 | -58036 | -15116 |
| LG+FQ | -53210 | -52504 | -15492 |
| LG+FO | -52325 | -51407 | -15643 |
| LG+FO+G12 | -49123 | -49071 | -16944 |
| GTR20+FO+G12 | -49054 | -49012 | -16930 |
| Simulated Abl/Src-Kinase (ESR) | | | |
| Model | true *LnP* | e*LnP* | SMP *LnP* |
| Poisson+FQ | -78347 | -76697 | -22585 |
| LG+FQ | -70969 | -69782 | -23505 |
| LG+FO | -69794 | -68587 | -23699 |
| LG+FO+G12 | -64530 | -64605 | -26327 |
| GTR20+FO+G12 | -64439 | -64513 | -26337 |
| Simulated Terpene Synthase (ESR) | | | |
| Model | true *LnP* | *eLnP* | SMP *LnP* |
| Poisson+FQ | -576140 | -567741 | -196181 |
| LG+FQ | -532336 | -525463 | -210825 |
| LG+FO | -523503 | -517089 | -211090 |
| LG+FO+G12 | -490634 | -490556 | -229746 |
| GTR20+FO+G12 | -490535 | -490394 | -229740 |

SI Table 11: Summary of total *LnP* as in Supplementary Table 7, but with extant reconstructions from data simulated using the LG+FO+G12 model of evolution from Supplementary Figures 9*c*-*e*, 10*c*-*e* and 11*c*-*e*
